# Secreted small RNAs of *Naegleria fowleri* are biomarkers for diagnosis of primary amoebic meningoencephalitis

**DOI:** 10.1101/2025.01.11.632551

**Authors:** A. Cassiopeia Russell, Joseph Dainis, Jose Alexander, Ibne Karim M. Ali, Dennis E. Kyle

## Abstract

Rapid and accurate diagnostics are needed to effectively detect and treat primary amoebic meningoencephalitis (PAM) caused by the free-living amoeba, *Naegleria fowleri*. Delayed diagnosis and similarities to other causes of meningitis contribute to a case mortality rate of >97%, and current testing requires a spinal tap to obtain cerebrospinal fluid (CSF). Thus, there is an unmet medical need for a non-invasive liquid biopsy diagnostic method. We sequenced *N. fowleri* extracellular vesicles (EVs) and identified microRNAs, tRNAs and other small RNAs in *N. fowleri-*EVs. From these data we selected two prevalent small RNAs as biomarker candidates. We developed an RT-qPCR assay and both small RNAs were detected in *N. fowleri-*EVs and amoeba-conditioned media. In the mouse model of PAM, both small RNA biomarkers were detected in 100% of mouse plasma samples at the end-stage of infection. Notably, smallRNA-1 was detected in the urine of infected mice at timepoints as early as 24h post infection (18/23 mice) and in the plasma as early as 60h post infection (8/8 mice). Additionally, smallRNA-1 was detected in 100% (n=6) of CSF samples from human PAM cases, and in whole blood samples, but not in human plasma from PAM cases. In this study, we discovered small RNAs as biomarkers of *N. fowleri* infection—one which can be detected reliably in CSF, urine, and whole blood. The RT-qPCR assay is a highly sensitive diagnostic assay that can be conducted in ∼3h after receipt of liquid biopsy. The data suggest detection of smallRNA-1 biomarker could provide earlier diagnosis of PAM and be used to monitor biomass of amoebae during treatment.

**One Sentence Summary:** Small RNAs of *Naegleria fowleri* can be detected in liquid biopsies and serve as early diagnostic biomarkers of primary amoebic meningoencephalitis.

## INTRODUCTION

Pathogenic free-living amoebae are found ubiquitously in soil and fresh water and cause highly lethal and under-diagnosed infections that pose significant risks to human health. *N. fowleri* only infects humans when amoeba-contaminated fresh water enters the nose, allowing the amoebae to invade the olfactory epithelium and traverse into the frontal lobes of brain to cause PAM(*1*). Though PAM remains a rare disease, with the Centers for Disease Control and Prevention (CDC) reporting 0-8 laboratory-confirmed cases in the U.S. annually(*2*), it is almost universally fatal with a mortality rate of >97%(*3*). Additionally, the geographic distribution of PAM cases in the U.S. and worldwide is expanding into more temperate regions in recent years—likely due to rising temperatures(*3–6*). Though our study focuses on *N. fowleri,* there are two other free-living amoebae that are known to cause disease in humans: *Acanthamoeba* spp., and *Balamuthia mandrillaris* (*7*)*. Acanthamoeba* spp. are opportunistic pathogens that predominantly infect immunocompromised individuals, 40% of patients infected with *B. mandrillaris* are immunocompromised, and *N. fowleri* mainly infects immunocompetent individuals(*8–10*). Though they are highly divergent phylogenetically(*11*), these amoebae are commonly reported together in the literature as all three can cause devastating neurological infections that are highly lethal(*8*). Due to the overlap in neurological symptoms associated with central nervous system infections caused by these pathogenic free-living amoebae, multiple drug discovery studies(*12–14*) and diagnostic development efforts(*15, 16*) have identified potential treatments and developed multiplexed assays targeting all three amoebae in the CSF. There remains a need, however, for diagnostic techniques that involve less invasive methods of obtaining biofluids compared with CSF to decrease the mortality related to acute PAM infections(*3*).

Current PAM diagnostic techniques rely almost entirely on detection in CSF. Methods include identification of motile trophozoites in wet mounts, visualizing amoeba with Wright-Giemsa staining, or sending CSF to an external reference facility for real-time PCR targeting *N. fowleri* DNA(*1*). Techniques other than PCR have several weaknesses, including the fact that amoebae closely resemble macrophages in appearance and motility and are often overlooked when microscopically analyzing CSF samples(*17*). Additionally, in multiple case reports, motile amoebae were not identified with the first and sometimes, even a second lumbar puncture. When diagnosing PAM infections via real-time PCR of CSF, the time that it takes to collect CSF, send samples to external diagnostic centers, and then wait for results is not optimal for such an aggressive, fulminant disease. In fact, PAM diagnoses are often confirmed postmortem or within 24 h prior to patient death(*18–21*). Recently we and others have characterized EVs that are secreted by *N. fowleri* (*22–25*) and we hypothesized that cargo in the EVs could serve as biomarkers of *N. fowleri*-infection. Herein, we identify secreted biomarkers for PAM infection that can be detected in less invasively collected biofluids and provide an RT-qPCR assay that can be performed with a thermal cycler and a qPCR machine within hours after obtaining the sample.

## RESULTS

The pipeline used to identify microRNAs, tRNAs and small RNAs secreted by *N. fowleri,* and a subset of the most prevalent predictions is presented in Fig. 1A-D (complete data in Supplemental Materials Table S1). A comparison between the top microRNAs identified in intact Nf69 trophozoites to those identified in Nf69-secreted EVs is shown in Supplemental Fig. S1. RT-qPCR assays confirmed the presence of three microRNAs in intact trophozoites of seven different clinical isolates (Supplemental Table S2 and Fig. S2), and two tRNAs in *N. fowleri* trophozoites (Villa Jose strain (VJ)), Nf69 EVs, and in the plasma of an Nf69-infected mouse (Supplemental Fig. S3). Assays were designed and standard curves were generated to target two highly prevalent biomarker candidates: smallRNA-1 and smallRNA-2 (Fig. 2A and C). Secretion of these small RNAs was confirmed by testing the EVs extracted from as many as 7 *N. fowleri* clinical isolates (Fig. 2A-D). To determine the specificity of the smallRNA-1 assay, the EVs of other pathogenic free-living amoebae (*A. castellanii* and *B. mandrillaris*) as well as 2 non-pathogenic *Naegleria* spp. were tested. The results indicated that this small RNA is potentially a *Naegleria* genus-specific biomarker as it was present in *N. lovaniensis* EVs and produced a slightly larger PCR product in *N. gruberi* EVs (Fig. 2E; gel confirmation and sequence alignment in Supplemental Fig. S4). Importantly, smallRNA-1 was not detected in *A. castellanii* or *B. mandrillaris* EVs.

**Fig. 1:**
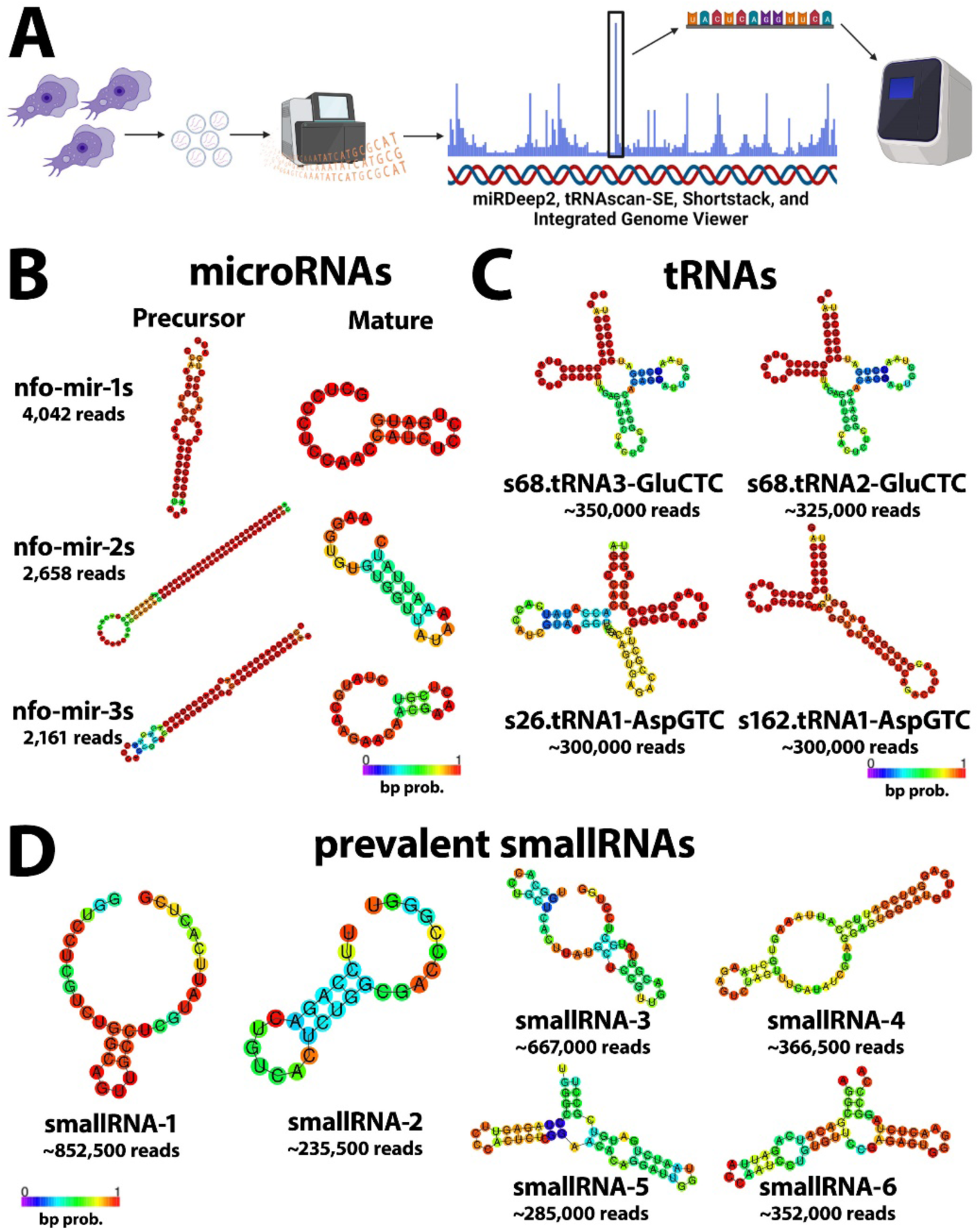
*Naegleria fowleri* encodes a diverse repertoire of small RNAs. (A) Schematic showing *Naegleria fowleri-*secreted EV small RNA identification pipeline. (B) Top 3 most prevalent precursor and mature microRNAs identified in whole cell small RNA sequencing with miRDeep2. (C) Top 4 most prevalent tRNAs identified in secreted EV small RNA sequencing with tRNAscan-SE. (D) Top 6 highly prevalent small RNAs in secreted EV RNAs identified manually via read-stacking with IGV. SmallRNA-5 is a tRNA fragment of s68.tRNA2-Glu. The 2 most prevalent small RNAs were identified first with Shortstack and confirmed manually with IGV. RNA secondary structures were generated with RNAfold v2.5.1(*33*). Schematic in panel A was generated with BioRender (www.biorender.com).

**Fig. 2:**
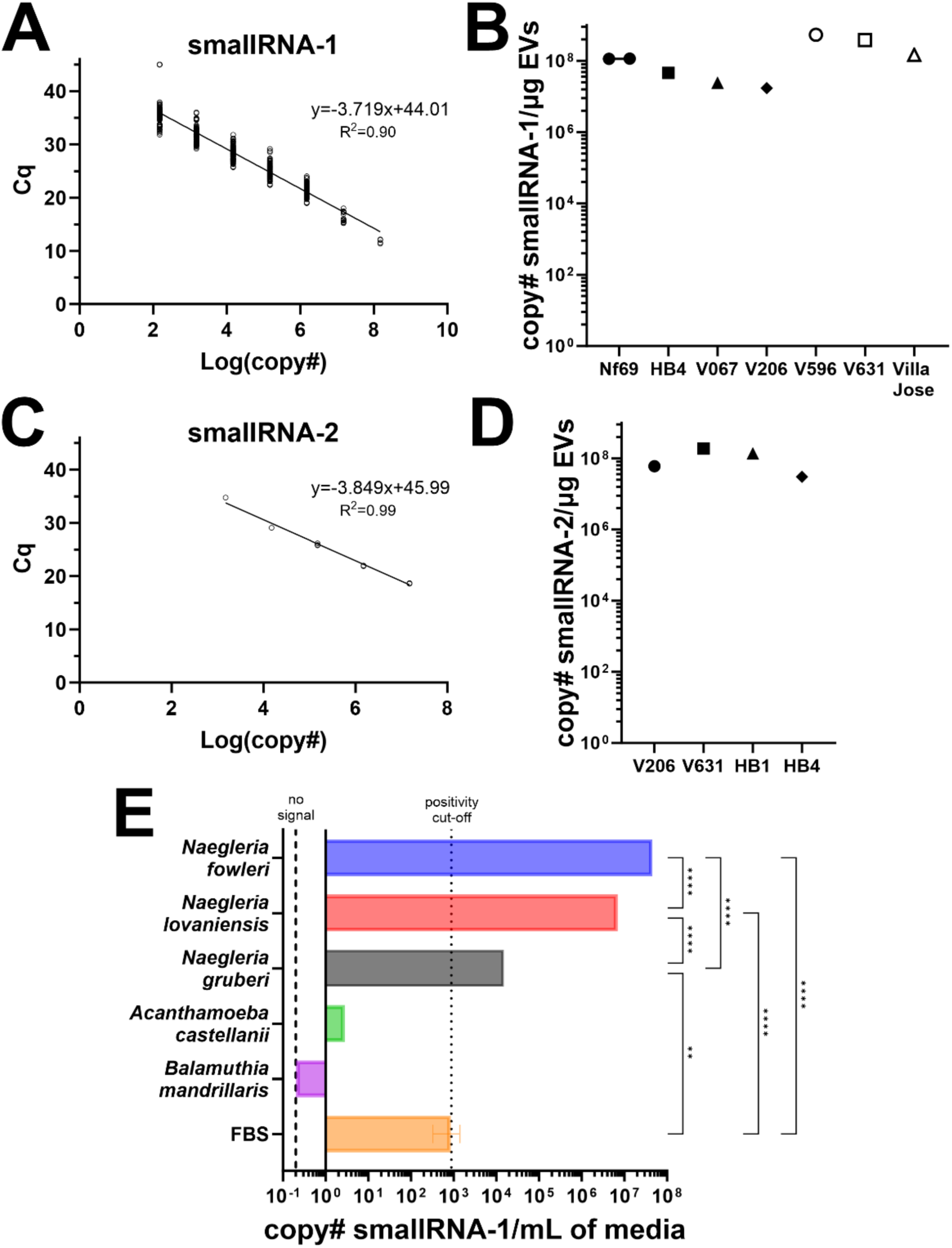
Detection of smallRNA-1 and smallRNA-2 in the EVs of various pathogenic and non-pathogenic free-living amoebae. (A) Standard curve for smallRNA-1 assay (consisting of 675 standards across 43 qPCR plates) was used to calculate copy number for qPCR reactions. (B) smallRNA-1 detection in RNA of EVs extracted via ultracentrifugation across 7 different clinical isolates of *N. fowleri*. Cq values ranged from 10.5-16.3. (C) Standard curve for smallRNA-2 assay used to calculate copy number. (D) smallRNA-2 detection in RNA of EVs extracted via ultracentrifugation across 4 different isolates of *N. fowleri*. Cq values ranged from 13.2-16.3. (E) smallRNA-1 detection in RNA of amoebae EVs extracted via extraction reagent from various species of free-living amoebae. For FBS and non-*Naegleria* species Cq values ranged from 33 to no signal. For *Naegleria* species Cq values ranged from 14.1-27.4. Each data point and/or bar in panels B-D consists of 3 technical replicates in RT-qPCR assay. Statistical significance was determined for panel E using One-way ANOVA test in GraphPad Prism v10.0.0 (GraphPad, La Jolla, CA, USA).

By testing RNA extracted from dilution series of conditioned media preparations in 96-well plates, we determined that the limit of detection for the smallRNA-1 assay is 10 to 100 amoebae cultured in 100µL of media for 24h (Fig. 3A); conversely, when we directly extracted from serial dilutions of *N. fowleri* trophozoites, we determined an limit of detection of <1 amoeba (Fig. 3B). Extraction and testing of conditioned media from wells containing axenic amoebae or amoebae feeding over Vero cells (Fig. 3C) indicates that smallRNA-1 is potentially secreted at higher levels while amoebae are feeding on mammalian cells versus while growing axenically (Fig. 3D). To ascertain translatability of the smallRNA-1 and -2 assays to *in vivo* settings, we tested the plasma of mice infected by 6 *N. fowleri* clinical isolates collected at the end-stage of infection (Fig. 4A) and obtained 100% positivity compared to uninfected human and mouse plasma for both small RNAs (Fig. 4B-C). We then performed a time course mouse infection study with the VJ isolate of *N. fowleri* (Fig. 5A; RNA extraction efficiency determined via spike-in in Supplemental Fig. S5; mouse survival curve and weight tracking in Supplemental Fig. S6A-B) and determined that smallRNA-1 can be detected in the serum at end-stage of disease, in the plasma as early as 60h post infection, and in the urine as early as 24h post infection (Fig. 5B; individual animal urine tracking in Supplemental Fig. S6C). To further ascertain the reliability of urine as a target biofluid, we infected an additional cohort of 10 mice with 5,000 Nf69 amoebae (RNA extraction efficiency, survival curve and weight tracking in Supplemental Fig. S5B, 6D and 6E, respectively) and obtained 100% positivity in the urine at ≥24h post-infection (Fig. 5C; individual urine tracking in Supplemental Fig. S6F).

**Fig. 3:**
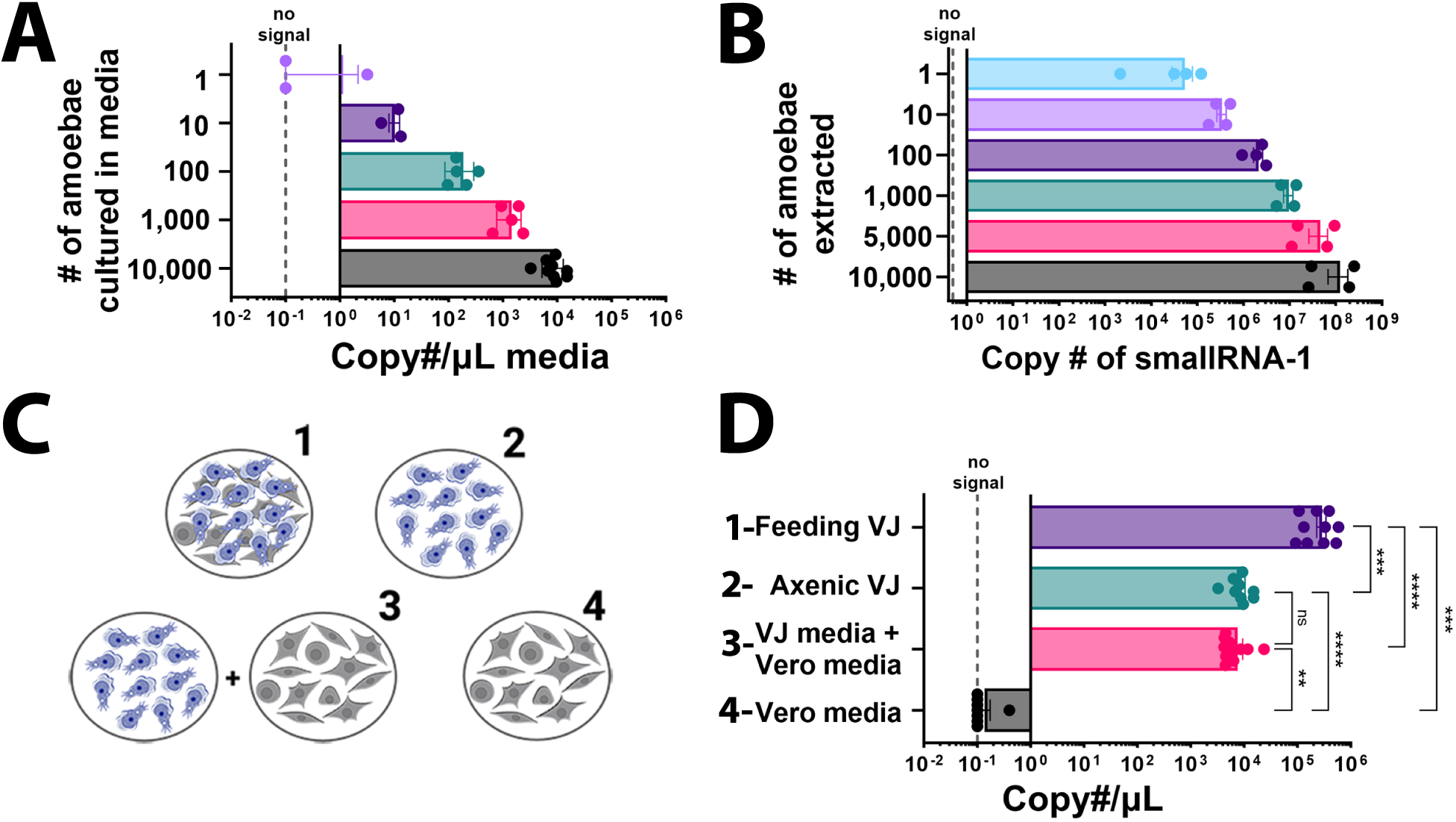
Detection of smallRNA-1 via RT-qPCR in *N. fowleri* media (A), intact trophozoites (B), and media with amoebae alone or feeding on Vero cells (D). (A) *N. fowleri* Villa Jose strain (VJ) amoebae were washed, diluted to final concentrations (1 – 10,000 amoebae), and cultured for 24h. Then cell suspensions were centrifuged and media extracted for quantitation of smallRNA-1. (Cq range: 24.8 to no signal.) (B) Trophozoites of N. fowleri were washed, centrifuged and RNA was extracted from amoebae pellets. smallRNA-1 was detected in amoebae diluted from 10,000 to 1 amoeba (Cq range for 1 amoeba: 29-35.5). (C) Schematic showing experimental set-up for assay to assess detection of smallRNA-1 in media from amoebae cultured (alone), from Vero cells (alone), and in cultures of amoebae feeding on Vero cells. Mixture of media from amoebae and Vero cultured alone were used as a control to assess if Vero cell-conditioned media affected the smallRNA-1 quantification. For each condition cells were incubated for 24 h before RNA extraction. (D) smallRNA-1 levels detected in media extracted from wells cultured for 24h (as shown in C) containing: 1-VJ feeding on Vero cell monolayer (Cq range: 19-22), 2-axenically cultured VJ (Cq range: 24.9-27.4), 3-axenic VJ media mixed with Vero media (Cq range: 23.1-25.9), 4-Vero media (Cq range: 41.8 to no signal). Each data point in panels A, B and D represents a single well in a 96-well plate, or a single amoeba pellet with 3 technical replicates in RT-qPCR assay. Statistical significance was determined for panel D using Unpaired T tests in GraphPad Prism v10.0.0 (GraphPad, La Jolla, CA, USA). Schematic in panel C was generated with BioRender (www.biorender.com).

**Fig. 4:**
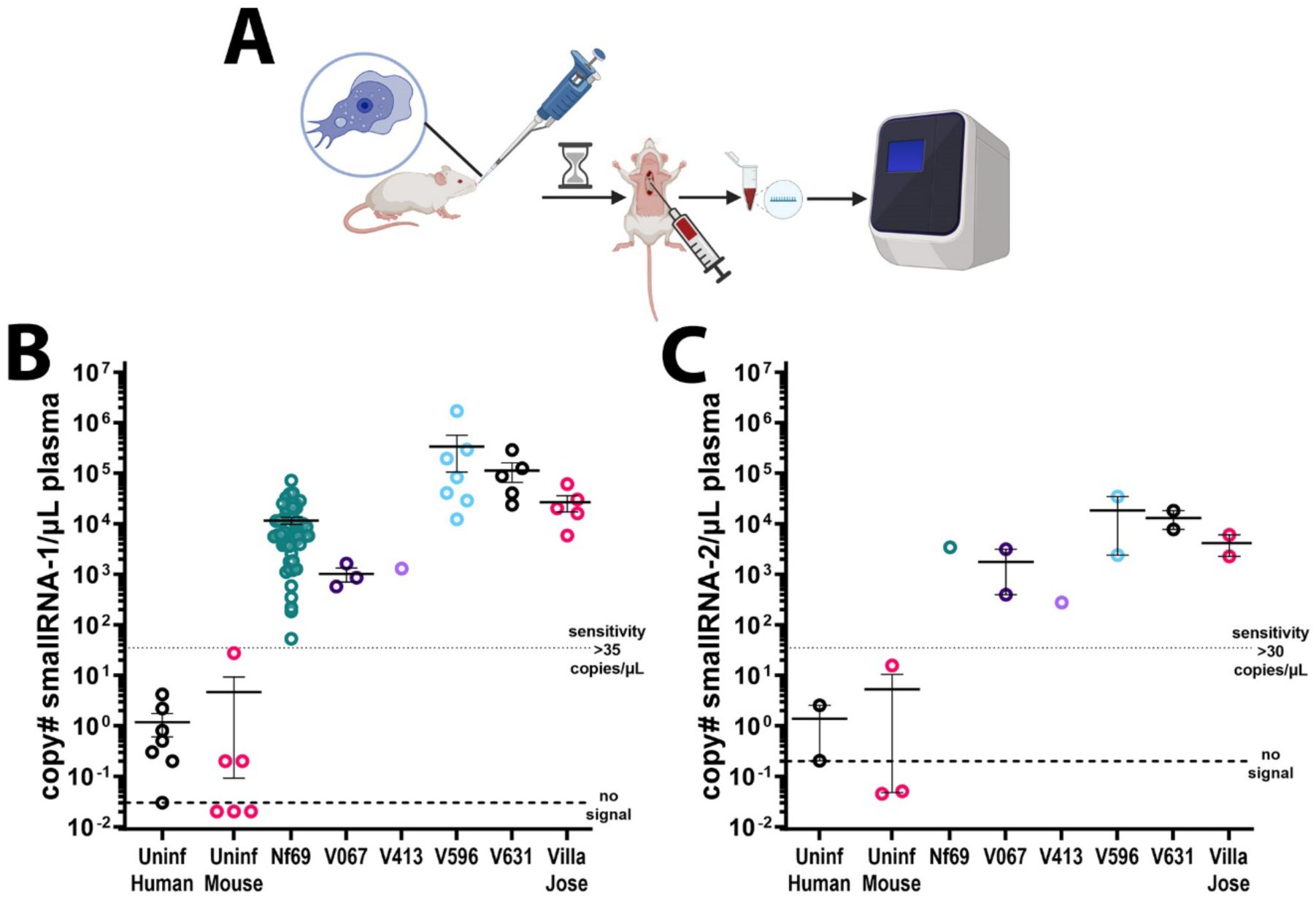
Detection of smallRNA-1 and -2 in the plasma of PAM-infected mice at the end-stage of infection. (A) Schematic showing infection process, plasma extraction via cardiac puncture followed by RNA extraction and RT-qPCR. (B) Assaying for smallRNA-1 in plasma of mice infected by 6 different *N. fowleri* clinical isolates provided 100% positivity in our assay compared to the uninfected mouse and human plasma. (C) Assaying for smallRNA-2 provided similar results to smallRNA-1 albeit at slightly lower concentrations per µL of plasma. All *N. fowleri*-infected mice were confirmed positive for amoebae by culturing brains and observing amoebae. Each data point in panels B-D is representative of 3 technical replicates in RT-qPCR assay. Schematic in panel A was generated with BioRender (www.biorender.com).

**Fig. 5:**
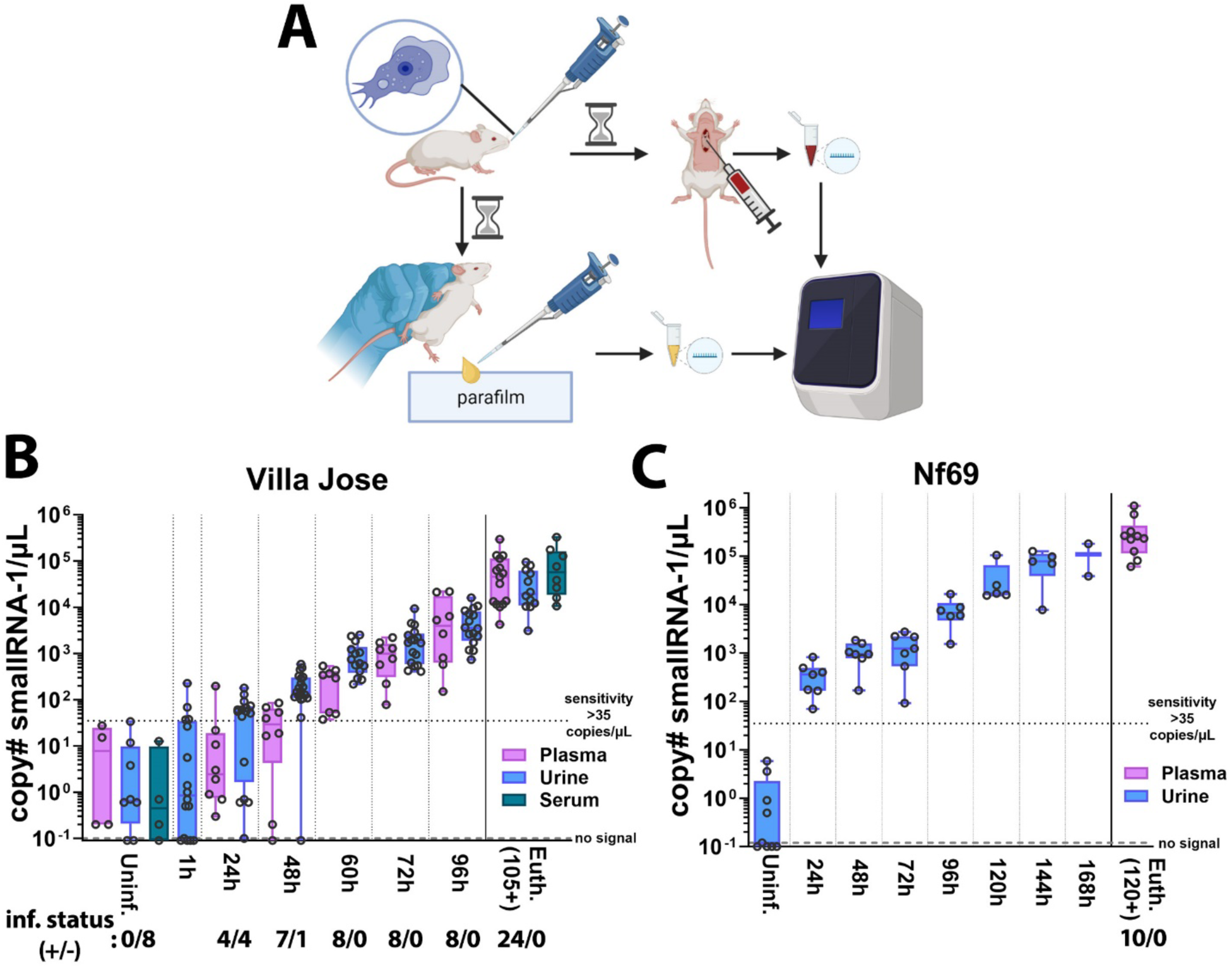
Detection of smallRNA-1 in the plasma, serum, and urine of PAM-infected mice at various timepoints post-infection. (A) Schematic showing infection process followed by urine collection at various timepoints and blood collection at the end-stage of infection followed by RNA extraction and RT-qPCR. (B) A cohort of 64 mice was infected with 1,000 VJ amoebae (fed over Vero cells 7 times) and cohorts of 8 mice each were sacrificed at various timepoints to extract blood. Urine was extracted from mice throughout infection. Mean time to death was 116.6h post infection. (C) A cohort of 10 mice was infected with 5,000 Nf69 amoebae (fed over Vero cells 8 times) and urine was either collected before infection or at various timepoints post-infection with plasma being extracted with euthanasia. Mean time to death was 140h post infection. The infection status of each cohort for panels B-C was determined by culture positivity of mouse brains and is shown under graphs. Each data point in panels B-C is representative of 3 technical replicates in RT-qPCR assay. “Positive” signals (>35 copies/µL) were defined as Cq values of 19.2 to 35.1 for panel B and 17.1 to 35.1 for panel C. Schematic in panel A was generated with BioRender (www.biorender.com).

To determine the potential of the assay for detecting *N. fowleri* in human samples, uninfected CSF, plasma, serum, and urine were tested to calculate a mean Cq positivity cut-off of <33.4 cycles (Fig. 6A and Supplemental Table S5). The limit of detection in human CSF, plasma, and urine was determined to be ∼100 copies/µL by spiking known dilutions of synthetic smallRNA-1 into 200µL aliquots of uninfected biofluids. Testing also indicated that serum sporadically inhibits both the smallRNA-1 and spiked-in cel-mir-39 RT-qPCR assays (Fig. 6A; RNA extraction efficiency in Supplemental Fig. S5B). Additionally, stability testing of Nf69 EVs spiked into CSF, plasma and urine indicates that EVs and their contents appear to degrade when stored at room temperature compared to 4°C (Supplemental Fig. S7). For further assay validation, CDC provided 15µL aliquots of CSF from confirmed *N. fowleri*, *B*. *mandrillaris*, and *Acanthamoeba*-infected individuals (Fig. 6B and Table S9). smallRNA-1 was detected in all *N. fowleri*-infected samples, whereas all CSF samples from *B. mandrillaris*- and *Acanthamoeba*-infected individuals were negative. For proof-of-concept in blood and serum, two *N. fowleri*-infected whole blood samples (20µL) and one plasma sample were tested. Notably, one of the whole blood samples as well as the single plasma sample were obtained from a survivor without an active infection. Results were positive for both whole blood samples, but the single convalescent plasma sample from the survivor was negative (Fig. 6C and Supplemental Table S9).

**Fig. 6:**
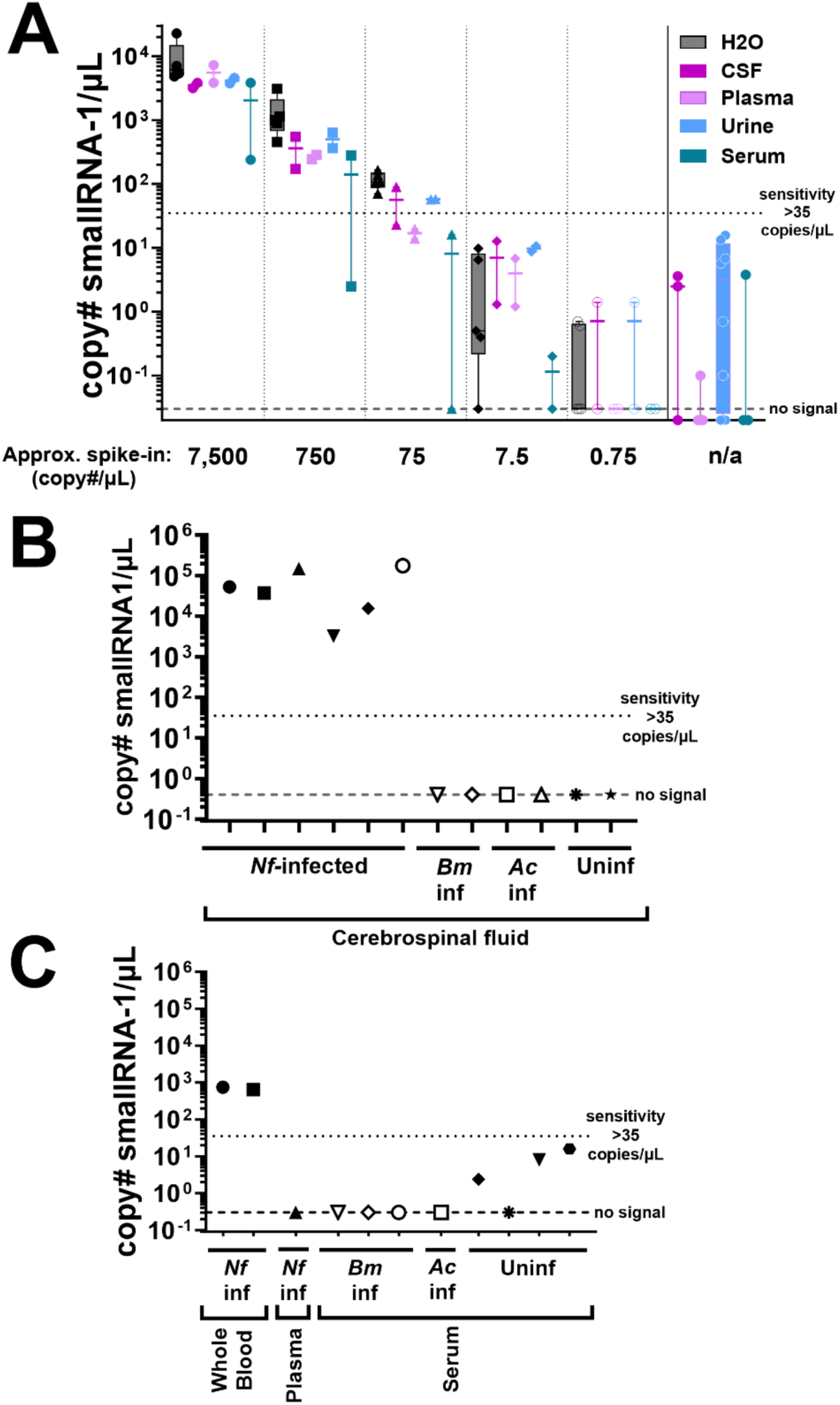
Determination of sensitivity of smallRNA-1 assay in various human biofluids compared to water and detection in PAM-infected human cerebrospinal fluid and whole blood. (A) SmallRNA-1 spike-ins into various biofluids indicate that CSF, plasma, and urine provide the most consistent results compared to H_2_O spike-ins, while serum seems to inhibit the assay. CSF, plasma, and serum were pooled from multiple individuals, whereas urine was tested from 8 individual samples. All human biofluid samples were de-identified and provided by AdventHealth Hospital, Orlando, Florida. (B) Detection of smallRNA-1 in PAM-infected (n=6), *Acanthamoeba*-infected (n=2), *B. mandrillaris-*infected (n=2), and uninfected human CSF (n=2) (Cq range for PAM-infected CSF: 24-30.8). (C) Detection of smallRNA-1 in *N. fowleri*-infected, uninfected, *B. mandrillaris*-infected, and *Acanthamoeba* spp.-infected human whole blood, convalescent plasma, and serum (Cq range for PAM-infected whole blood: 32.4-32.6). We adopted a “positivity” cut-off of <33 cycles for urine, and <35 cycles for other biofluids; equivalent to > ∼100 copies per µL of biofluid). Each data point is representative of 3 technical replicates in RT-qPCR assay.

## DISCUSSION

This study provides the first evidence that the EV-secreted small RNAs of *N. fowleri* are biomarkers that can be detected with RT-qPCR assays in multiple biofluids, including CSF and whole blood for humans, and urine and plasma for mice. PCR of *N. fowleri* DNA in CSF is the current golden standard for PAM diagnosis. Alternatively, or in addition to, amoebae can be identified microscopically in CSF. These methods require collection of CSF and most often amoebae are detected postmortem or too late in infection for current drugs to improve clinical outcomes. One study detected *N. fowleri* infection in the blood of a deceased patient by using meta-genomic next-generation sequencing (*26*); however, this method is not ideal for PAM diagnosis since it requires several days between sample collection and analysis of sequence data.

In this study we identified *N. fowleri* smallRNA-1 as a reliable biomarker of infection and propose that an RT-qPCR assay could be used for much earlier diagnosis of infection with *N. fowleri.* Given the aggressive nature of PAM and the current difficulties to differentially diagnose amoebae versus bacterial or viral meningitides, the new biomarker offers clinically significant advantages that could improve on the current ∼97% case fatality rate for PAM. The smallRNA-1 early-stage biomarker is also advantageous as it is detectable in fluids that can be obtained through less invasive procedures than the currently used qPCR diagnostic assays. The biomarker assay also offers the potential for monitoring the biomass of amoebae via urine or blood sampling during treatment. At present there are no accurate measures of amoebae density that can assist physicians with timely monitoring of treatment efficacy. Potentially, combining multiple small RNAs identified in this study could improve the already significant sensitivity and specificity of the RT-qPCR assay for smallRNA-1 alone. RT-qPCR based diagnostics are becoming more widely adopted in clinical centers following the recent SARS-CoV-2 pandemic(*27*). This development provides the capability of in-house RT-qPCR assays of biofluids extracted in point-of-care situations. Further clinical adaptations to point-of-care testing such as transitioning to a one-step RT-qPCR assay to reduce complexity and time or testing nasal swabs/saliva could also increase the utility of this assay.

The data from infecting mice with *N. fowleri*, spiking smallRNA-1into human biofluids, and testing biofluid samples from PAM patients suggest multiple biofluids could be used to detect the biomarker. CSF, urine, whole blood, and plasma provided the most consistent results, whereas apparent degradation was observed in serum (Figure 6). The only plasma specimen available from a confirmed PAM patient was negative for smallRNA-1 (Figure 6C); however, it was collected from a survivor one month after admission to hospital. The patient had fully recovered before the convalescent plasma was collected. Therefore, our interpretation is the active infection with amoebae was cleared and the biomarker did not persist in circulation. Additional studies are warranted with more human plasma samples to confirm this result and to establish if the biomarker is present only during an active infection.

One limitation of this study is the small sample size of biofluids from *N. fowleri*-infected patients that could be tested. Due to the rarity of this disease, it is unlikely that large numbers of specimens from infected humans can be collected to further validate the biomarker. Additional animal studies could be substituted for this purpose. We also found that smallRNA-1 appears to be a *Naegleria* genus specific marker; however, given that liquid biopsies for this assay can be collected aseptically, the potential for false positives from naturally occurring *Naegleria* spp. can be obviated. Regardless of any advances made in the realms of diagnosis and treatment, it is critical that attending medical professionals are familiar with free-living amoebae infections including PAM, that suspicion is raised if there is a history of warm freshwater exposure, and that early differential diagnoses include free-living amoebae as potential etiologic agents. More detailed guidance for health care providers regarding clinical features, diagnosis, and treatment can be found at the website provided by US Centers for Disease Control (https://www.cdc.gov/parasites/naegleria/health_professionals.html).

## MATERIALS AND METHODS

### Amoeba Culturing

*N. fowleri* clinical isolates and *N. lovaniensis* were cultured as previously described(*28*), with some minor modifications. Amoebae were either grown axenically or were extracted from infected mice by placing infected brain in Nelson’s complete media (NCM) supplemented with 10% fetal bovine serum (FBS; Corning, Oneonta, NY, USA) and 1000 U/mL penicillin and 1000mg/mL streptomycin (Gibco, Gaithersburg, MD, USA) in non-vented 75-cm^2^ tissue culture flasks (Olympus, El Cajon, CA, USA) at 34°C and 5% CO_2_. To confirm *N. fowleri* mouse infection, extracted brains were incubated in flasks until adherent amoebae were visualized and pure amoebae cultures from the infected brains were established. *N. gruberi* was cultured in M7 media(*29*) supplemented with 10% FBS at 27°C in in non-vented 75-cm^2^ tissue culture flasks. *B. mandrillaris* and *Ac* were cultured as previously described(*12*). Amoebae were passaged by placing *Naegleria* spp. flasks on ice for 10-15min followed by light mechanical tapping for *N. gruberi* to detach cells, by mechanically harvesting *Ac* from flasks(*30*), or by incubating *B. mandrillaris* with 0.25% Trypsin-EDTA (Gibco) for 5min at 37°C to detach cells from flask. The resulting cell suspensions were centrifuged at 3,900rpm for 5min at RT. A hemocytometer was used for counting amoebae in duplicate for reported cell concentrations.

### Nf69-Conditioned Media EV Extraction for RNA sequencing

Nf69 was grown from an infected mouse brain in a non-vented 75-cm^2^ tissue culture flask and washed several times with 1X phosphate buffered saline (1XPBS; Gibco) to obtain a confluent amoeba culture. Contents of flask were iced for 10-15min and transferred to a 175-cm^2^ tissue culture flask with a media change and 1XPBS wash once amoebae reached logarithmic stage. Media was replaced with NCM supplemented with 10% EV-depleted FBS (Gibco) and amoebae were allowed to adapt to new media for a week with media changes followed by PBS washes to remove cell debris until amoebae were growing robustly. A final volume of 100mL of media supplemented with 10% EV-depleted FBS and 10% penicillin/streptomycin was added and amoebae were allowed to grow to confluency prior to harvesting cells and conditioned media by placing on ice for 15-30min and centrifuging at 3,900rpm for 15min at 4°C. Extraction details for sample 1 (passage 0) and sample 2 (passage 1; passed from sample 1) are shown in Table S4.

Conditioned media was transferred to new tubes without disturbing cell pellets which were saved for counting in duplicate. Total Exosome Isolation (from cell culture media) reagent (Invitrogen, Waltham, MA, USA) was used according to manufacturer protocols to extract EVs from amoeba conditioned media. Resulting EV extractions were resuspended in 4mL of 0.1µm-filtered 1XPBS each. These suspensions were stored at -20°C until being sent on dry ice for small RNA sequencing to System Biosciences (SBI; Palo Alto, CA, USA) for their Exo-NGS Exosomal RNA Sequencing services. Raw data was submitted to the National Center for Biotechnology Information’s (NCBI) Sequence Read Archive (SRA) and can be found under BioProject ID PRJNA991274. Pre-processing protocols are available in Supplemental Materials.

### Prevalent small RNA identification via Shortstack and Integrated Genome Viewer

Shortstack v3.8.5(*31*) was used to identify small RNA clusters with dicermax option set to 100 and sort-mem set to 1000M. The MajorRNAs within the identified clusters with the most reads were manually confirmed by importing small RNA alignment and Shortstack output .gff files into Integrated Genome Viewer v2.8.10 to visualize read-stacking peaks against the Nf69 genome (BioProject ID PRJNA1002350) and sequences for top/most prevalent peaks were extracted for qPCR targeting and confirmation (provided in Table S1). All Shortstack output files are provided in the Mendeley data repository.

### RNA extractions and cel-mir-39 spike-in for RNA extraction efficiency evaluation

For small RNA extractions from whole amoebae, EV suspensions, conditioned media, mouse biofluids and human biofluids, the mirVana miRNA isolation kit (Invitrogen) was used according to the manufacturer’s protocols with alterations described below. For whole amoebae, amoebae were first counted in duplicate, transferred to microcentrifuge tubes, centrifuged for 3min at 14,000rpm, and supernatant was discarded before adding 300µL of Lysis/Binding buffer to cell pellet. For cel-mir-39 spike-ins, the microRNA (cel-mir-39) Spike-In Kit (Norgen Biotek Corp., Thorold, Ontario, Canada) was used to measure RNA extraction efficiency, 3µL of the provided 33fmol cel-mir-39 suspension was diluted in a final volume of 56µL of Nuclease-Free Water to mimic the elution volume of 56µL used for RNA extractions. For spiking into fluids, 3µL cel-mir-39 was added to lysis solution prior to adding 1/10^th^ volume of miRNA homogenate additive and proceeding with manufacturer’s protocols. For RNA extractions from media and other fluids, Lysis/Binding buffer equivalent to 2x volumes of biofluid/media was added. If less than 150µL of fluid was used for extraction, a minimum volume of 300µL of Lysis/Binding buffer was added before following the process described above. Samples were eluted with 56-100µL of Elution Solution that was preheated (final optimized protocol uses 56µL elution volume to obtain more concentrated RNA suspension) to 95°C and stored at -80°C until use in RT-qPCR assay.

### Conditioned Media Preparation and EV extractions for small RNA detection

To generate conditioned media for EV extractions, amoebae were cultured and EVs were extracted and protein concentrations were measured as previously described(*22*). To extract EVs from other free-living amoebae, we cultured them in their respective culture media and used the previously described Total Exosome Isolation reagent to extract EVs from 25mL of conditioned media (see Supplemental Table S3). To avoid the process of adapting each species to EV-depleted FBS, we also extracted EVs from 2.5mL of fetal bovine serum to mimic the 10% supplementation in media for *N. fowleri*, *B. mandrillaris*, *N. lovaniensis* and *N. gruberi* (*A. castellanii* media contains no FBS).

### Taqman small RNA RT-qPCR assays

Custom Taqman Small RNA Assays (Applied Biosystems, Waltham, MA, USA) were designed by submitting the target sequences to the Custom Small RNA Design Tool with resulting assay IDs and cycling parameters for all Taqman assays used in this study provided in Supplemental Tables S2 and S3. RNAs were reverse-transcribed in 96-well PCR plates (Thermo Scientific, Waltham, MA, USA) using the TaqMan® MicroRNA Reverse Transcription kit (Applied Biosystems, Waltham, MA, USA) by first creating RT Reaction Mix and placing on ice, and then combining 5µL of total RNA elution volume and 3µL of the RNA–specific stem–looped RT primer and proceeding with the initial pre-processing step recommended by manufacturer for double-stranded small RNA. Reverse transcription was then performed according to manufacturer protocols. A no reverse transcriptase and a non-template RT control was included in each reaction plate. qPCR was performed with TaqMan Fast Advanced Master Mix (Applied Biosystems) according to manufacturer’s protocols—with the alteration of using 18µL of PCR master mix with a volume of 2µL cDNA from reverse transcription reaction—using a CFX96 Real-Time System/C1000 Touch Thermal Cycler. Three non-template PCR controls (one for cel-mir-39) were included in each reaction plate. Each sample was assayed in triplicate (duplicate for cel-mir-39) and the mean Cq value was used to calculate the copy number. To generate standard curves, RNase-Free HPLC purified synthetic oligos were ordered from Integrated DNA Technologies (Coralville, IA, USA) and resuspended to 10uM followed by dilutions and calculations according to the techniques described by Kramer et al(*32*), with the alteration of 2µL cDNA input to qPCR rather than 1.3µL to increase sensitivity.

### 96-well plate culture media assay

Vero cells (green monkey kidney cells; E6; ATCC CRL-1586) were maintained as previously described(*22*), with the alteration of supplementing media with EV-depleted FBS. The night before amoebae were added, 25,000 Vero cells were seeded into wells of a 96-well tissue-culture treated plate to generate a full monolayer. The next morning, media was replaced with fresh pre-warmed media and wells were either left as controls or seeded with 10,000 Villa Jose amoebae. Additionally, ranges of concentrations of amoebae (1–10,000) were added to empty wells in EV-depleted NCM. These were allowed to grow/feed for 24h at 37°C and 5% CO_2_. Plates were then spun at 3,900rpm for 5min at 37°C and well supernatants were extracted into individual microcentrifuge tubes. These were then spun at 3,900rpm for another 5min at RT and the supernatants were placed into fresh tubes after which RNA extractions were performed as previously described.

### Animal Studies

All animal studies were performed in the laboratory animal facilities at the UGA College of Veterinary Medicine and were approved by the UGA Institutional Animal Care & Use Committee (Protocol A202 03-026). Animals were housed in conventional rooms, 22.2+/-2°C and 50+/-20% relative humidity with a controlled 12h-light/12h-dark cycle. The animals used in these studies were completely separated from other animals and maintained in accordance with the applicable portions of the Animal Welfare Act and the DHHS “Guide for the Care and Use of Laboratory Animals.” Veterinary care was under the supervision of a full-time resident veterinarian boarded by the American College of Laboratory Animal Medicine. Mice were euthanized by exposure to gaseous CO_2_ to induce narcosis and death, which is consistent with the recommendations made by the Panel on Euthanasia of the American Veterinary Medical Association. Prior to infecting mice with clinical isolates of *N. fowleri,* we first confirmed pathogenicity by transferring trophozoites from axenic culture conditions to flasks containing Vero monolayers, forcing the amoebae to actively feed. We passaged these amoebae populations over subsequent Vero monolayers 5-11 times prior to infecting anesthetized mice by inoculating 10µL amoeba suspensions into the right nare. Progression of infection was monitored with 3x daily observation of mice, with the development of pre-moribund signs or weight loss used as evidence of disease progression. Pre-moribund criteria include: hypoactivity, rapid and/or labored breathing, ataxia, soiled anogenital area, and/or hunched posture. If animals were found pre-moribund in these studies, they were euthanized. Body weight was measured daily, starting on the day of infection, and mice with a decrease in body weight of 20% or more (a sign of severe morbidity) were removed and euthanized.

In a pilot animal study, plasma was collected from female 3–4-week-old transgenic ICR (CD-1) mice that were infected by either a clinical isolate of *N. fowleri* (Nf69–10,000 amoebae n=52; V067–5,000 n=3; V413–20,000 n=1; V596–1,000 n=2, and 5,000 n=5; V631–5,000 n=5; Villa Jose–5,000 n=5), or *Plasmodium berghei* (n=6) as negative controls. Samples were randomly collected from mice infected with *N. fowleri* or *Plasmodium berghei*. Plasma containing CPDA-1 anticoagulant from human donors (n=7) was purchased from Grifols BioSupplies (Memphis, TN, USA) and used to test for signals in uninfected biofluids (details on these samples and other human biofluid controls are provided in Supplemental Table S5). RNA samples were tested initially for smallRNA-1 with the remainder of some being allocated for a proof-of-concept assay with smallRNA-2 and two tRNAs. For the first smallRNA-1 follow-up study, we planned a time course infection experiment consisting of 18 groups (1 group=1 cage of 4 mice) of 3-4-week-old ICR (CD-1) female mice (total n=72). This was undertaken by infecting 16 groups with 1,000 Villa Jose amoebae and keeping two uninfected control groups for plasma and urine collection to establish baseline signals for the assay. Two groups of mice (n=8) were euthanized at various timepoints post-infection (24-96h) to extract plasma and determine the earliest time post-infection that smallRNA-1 could be detected in the plasma. Throughout the study, urine was collected from mice from various infected groups with a similar goal. Additionally, serum was collected from two groups of end-stage infections in mice to confirm smallRNA-1 detection in this biofluid. Groups of mice were kept separate for the entirety of the study, and the order in which cages were pulled to extract urine from infected groups was randomized. The plasma for 1 out of 56 infected mice was excluded as this mouse succumbed to infection between the night check of day 6 and the morning check of day 7 post-infection and we were unable to obtain an extraction. For the final smallRNA-1 time course study, a smaller cohort (n=10) of two groups of 3-4-week-old ICR (CD-1) female mice (1 group=1 cage of 5 mice) were infected with 5,000 Nf69 amoebae and urine was collected daily with plasma collection post-euthanasia.

To collect urine, parafilm was placed in the bottom of biosafety cabinet and the lower abdomen of restrained mice was lightly depressed to induce urination onto the parafilm. Urine was collected with pipettors and transferred to microcentrifuge tubes prior to storage at -80°C within 1-2hr post collection until RNA extraction. To collect plasma post-euthanasia, 1mL syringes with 26g needles were primed with 50-100µL of ACD anticoagulant (Boston Bioproducts, Inc., Milford, MA; cat#: IBB-400) prior to sacrificing mice. Mice were then euthanized with CO_2_ and exsanguination via cardiac puncture was performed. Blood/anticoagulant suspension was then spun at 4,000rpm for 10min within 1-2h of collection. For serum, blood was extracted in the same manner but with no anticoagulant. Whole blood was allowed to coagulate for 45min-1h before spinning at 4,000rpm for 10min. The supernatant (plasma/anticoagulant or serum) was collected and frozen at -80°C until RNA extraction.

### Human Biofluid testing

Human biofluids (CSF, plasma (ACD anticoagulant), serum and urine), collected as part of the patient care at AdventHealth Orlando were deidentified and provided for spike-in testing. CSF, plasma and serum are representative of pooled samples while urine was not pooled. Deidentified, *N. fowleri* infected patient biofluids were provided by the Centers for Disease Control and Prevention after being stored at <-20°C for an undisclosed amount of time (also see Supplemental Table S9 notes).

## Data Sharing

All sequencing data is deposited to NCBI SRA under BioProject IDs PRJNA991265 and PRJNA991274. Extended data files are shared in Mendeley Data (doi: 10.17632/9psx79t365.1).

## Acknowledgments

The funder of the study played no role in study design, data collection, data analysis, data interpretation, or writing of the report. The findings and conclusions in this report are those of the authors and do not necessarily represent the official position of the U.S. Centers for Disease Control and Prevention.

## Funding

Georgia Research Alliance (DEK)

US National Institute of Allergy and Infectious Diseases T32AI060546 (ACR, DEK)

US National Institute of Allergy and Infectious Diseases R03AI141709 (DEK)

## Author contributions

Conceptualization: ACR, DEK

Methodology: ACR, JD, JA, IKMA, DEK

Investigation: ACR, JD

Visualization: ACR, JD, DEK

Funding acquisition: ACR, DEK

Project administration: DEK

Supervision: JA, IKMA, DEK

Writing – original draft: ACR, DEK

Writing – review & editing: ACR, JD, JA, IKMA, DEK

