## Supplemental Tables and Figures for "Secreted small RNAs of *Naegleria fowleri* are biomarkers for diagnosis of primary amoebic meningoencephalitis"

### Supplementary Methods

#### Amoeba Isolates

*N. fowleri* Nf69 (ATCC 30215) obtained from a 9-year-old boy in Adelaide, Australia in 1969, and *Acanthamoeba castellanii* type T4 (ATCC 50370) isolated from the eye of a patient in New York, NY in 1978 were purchased from the American Type Culture Collection (ATCC). *Balamuthia mandrillaris* (CDC:V039; ATCC 50209), a GAE isolate, isolated from a pregnant baboon at the San Diego Zoo in 1986 was donated by Luis Fernando Lares-Jiménez ITSON University, Mexico. *N. fowleri* Villa Jose, isolated from a female in California in 1996, V067, isolated from a 30-year-old male in Arizona in 1987, V413, isolated from a 17-year-old boy in Texas in 1998, V596, isolated from a male in Nevada in 2007, V631, isolated from a 28-year-old man in in Louisiana in 2011, and *N. lovaniensis* (ID: 76-15-250) [1] were all kindly provided by Dr. Ibne Ali at the Centers for Disease Control and Prevention. *N. gruberi* NEG-M (ATCC 30224) was kindly provided by Dr. Katrina Velle at the University of Massachusetts Amherst.

#### Sample Preparation and Small RNA Sequencing of whole *N. fowleri* amoebae

Two independent *Naegleria fowleri* trophozoite (Nf69; ATCC 30215) suspensions were counted, diluted, and spun at 14,000rpm for 3min prior to discarding supernatant, and RNA was extracted using a Direct-zol RNA miniprep kit (Zymo Research, Irvine, CA, USA; cat#: R2051) according to the manufacturer's protocol. Sample 1 consisted of $3x10^6$ trophozoites and sample 2 consisted of $5x10^6$ trophozoites. These two RNA samples were then sent to the University of Georgia Genomics and Bioinformatics Core for quality checking via an Agilent Bioanalyzer (results provided in Mendeley Data Repository) and small RNA library preparation with TruSeq small RNA library preparation kit (Illumina, Inc., San Diego, CA, USA), and subsequent sequencing using NEXTFLEX Small RNA-Seq Kit v3 (PerkinElmer, Austin, TX, USA) with Illumina NextSeq SE75 flow cells with four lanes per sample. Raw data was submitted to the NCBI SRA and can be found under BioProject ID PRJNA991265 with BioSample accessions SAMN36291057 and SAMN36291058.

#### Pre-processing of raw whole cell small RNA sequencing data for input to miRDeep2

The resulting whole cell small RNA sequencing data was pre-processed by trimming the sequencing adaptors with Cutadapt v2.8[2] according to the NEXTFLEX small RNA trimming recommendations, and 4 bps were also trimmed from either end as these randomized additions are specific to the NEXTFLEX kit. Following this, the data for the four lanes for each sample were merged into two files that were analyzed with FastQC v0.11.9 to confirm the efficacy of the adaptor trimming as well as the quality of the sequences. After confirming adequate quality, pseudo-ribodepletion was performed by using HISAT2 v2.1.0[3] to map the sequences to an rDNA plasmid (GenBank accession #: CM017919) for *N. fowleri* created by Leichti et al. [4] with the unmapped reads being extracted for downstream processing.

#### Pre-processing of raw EV small RNA sequencing data for input to miRDeep2

Raw FASTQ files for each sample were trimmed of adaptors with Cutadapt v2.8 using adaptor sequences 'TGGAATTCTCGGGTGCCAAGG' and

‘GATCGTCGGACTGTAGAACTCTGAAC’ and minimum length cut-off of 23 bp. These trimmed reads were then pseudo-ribodepleted in the same way that the whole cell small RNAs were, and we proceeded with reads that did not align to this plasmid. Post-trim FastQC v0.11.9 html reports as well as trimmed and ribodepleted EV sequencing data are provided in the Mendeley data repository.

**Pre-processing of raw EV small RNA sequencing data for input to Shortstack**

Raw FASTQ files for each sample were trimmed of adaptors in Galaxy version 23.1.rc1 (commit b58f5824abced2e090238000367e1ae1b5ef02df) with the software package Trim Galore v0.6.3 with Cutadapt v2.3 and Python v3.7.3 using the following options: quality Phred score cutoff=30, quality encoding type selected=ASCII+33, adapter sequence= ‘TGGAATTCTCGG’ (Illumina small RNA adaptor; auto-detected), maximum trimming error rate=0.1 (default), minimum required adapter overlap (stringency)= 1bp, minimum required sequence length before a sequence gets removed= 15bp. For sample 1, 22,006,563 reads were processed of which 98.4% (21,650,997) had adaptors and 100% passed filters. For sample 2, 18,077,331 reads were processed of which 99.8% (18,034,333) had adaptors and 100% passed filters. Pre- and post-trim FastQC html reports and trimmed EV sequencing data are provided in the Mendeley data repository.

**microRNA identification in whole cell small RNAs and EV RNAs via miRDeep2**

Pre-processed whole cell and *Nf*-EV small RNAs that did not align to the rDNA plasmid were first were run through the miRDeep2 v0.1.3[5] software package containing Bowtie module (which was used with Nf69 reference genome with default settings to build a reference index as required by the package), and mapper.pl module (which was also required by the package with options -e -h -i -j -l 18 -m). Resulting files processed by Bowtie and mapper.pl were input to main miRDeep2 module with ‘none’ input for mature reference miRNAs, ‘none’ for mature other miRNAs and ‘none’ for hairpin reference miRNAs. Reported microRNAs were confirmed in the sequencing results across both samples for the small RNAs sequenced from the whole cells and the EVs. The results were then cross-referenced to determine if microRNAs identified in the whole cell were also secreted in the EVs. All miRDeep2 output files are provided in the Mendeley data repository.

**tRNA identification via tRNAscan database**

The *N. fowleri* Nf69 reference genome (BioProject ID PRJNA1002350) was input to tRNAscan-SE v2.0.7[6] with search mode option -g for general tRNA model, --mid, and --notrunc to predict *N. fowleri* tRNAs. The output data was then manually compared to the EV-sequencing data aligned to the reference Nf69 genome within Integrated Genome Viewer to identify tRNAs secreted within the EVs based upon read-stacking (provided in Table S1 in Appendix). All tRNAscan-SE output files are provided in the Mendeley data repository.

**Additional small RNA identification methods**

The resulting output from ShortStack consisted of 9,913 clusters identified for sample 1 and 6,895 clusters identified for sample 2, resulting in an average of 8,404 clusters of small RNAs aligning to the *N. fowleri* Nf69 genome. When selecting most prevalent small RNAs, we utilized the primary criterion of highest number of reads per cluster followed by the secondary criterion of highest number of major RNA detected to select the top biomarker candidates in the ShortStack data. To audit discrepancies in sizes of the first major RNA predicted between the two samples (42 bp versus 34 bp), we utilized the Integrated Genome Viewer (IGV; v2.8.10) software to manually reconfirm the location of clusters aligning to the genome and to definitively select the small RNAs with the highest read stacking. Upon locating the clusters of aligned small RNA reads, we noted that reads of shorter lengths than those predicted by ShortStack seemed to be higher in prevalence, so we selected the overlapping sequence with the highest prevalence. This resulted in the same 34 bp sequence for smallRNA-1 predicted in sample 2 with ShortStack. We applied this same process to smallRNA-2 and upon locating the clusters of aligned small RNA reads, we found that reads of the same length and sequence as those predicted by ShortStack aligned to the genome within the predicted cluster. This allowed us to confidently move forward with two highly prevalent small RNA biomarker candidates.

**qPCR for nfo-mir-1, nfo-mir-2, and nfo-mir-3**

For qPCR confirmation of the first 3 microRNAs identified in whole amoeba small RNAs, we designed stem-loop reverse transcription primers according to the protocol supplied by Kramer[7] and provide the primer sequences, concentrations, and annealing temperatures in Table S2. Taqman microRNA Reverse Transcription kit was used according to manufacturer's protocols and qPCR was performed with Luna qPCR Master Mix (New England Biolabs, Ipswich, MA, USA) according to manufacturer's protocols,

**Agarose gel, cloning and sequencing of smallRNA-1 qPCR products**

To prepare smallRNA-1 samples, 1 technical replicate of qPCR products was cleaned and concentrated using a DNA Clean and Concentrator-5 kit (Zymo Research) and samples were eluted in 12µL of elution buffer. 3µL of concentrated qPCR products were combined with 2µL 6X Loading Dye and 7µL $H_2O$. These products and 10µL of Quick-Load Purple Low Molecular Weight DNA Ladder (New England Biolabs, Ipswich, MA, USA) were loaded and run on a 3% agarose gel stained with SYBR Safe DNA Gel Stain (Invitrogen, Waltham, MA, USA) for 2h at 80V.

We then inserted the concentrated qPCR products into vectors using a CloneJET PCR Cloning Kit (Thermo Fisher Scientific, Denver, CO, USA) and cloned these into the provided DH10B competent cells according to manufacturer's protocols for both blunt and sticky ends. We performed blue/white colony screening and amplified several replicates of white colonies overnight before performing minipreps using a Quick-DNA Miniprep Plus kit (Zymo Research). We confirmed the insertion of the qPCR product with PCR of miniprepped plasmids using the AccuPrime Taq DNA Polymerase System (Invitrogen) according to manufacturer's protocol using the amplification primers provided by the CloneJET PCR Cloning Kit (Thermo Scientific). After running on an agarose gel, positive clones were confirmed by visualization of an increase in band size relative to negative clones. The resulting confirmed plasmids were sent to Genewiz (South Plainfield, NJ, USA) for Sanger sequencing in both the forward and reverse directions using the provided pJET1.2 sequencing primers. Poor-quality bases were trimmed from either end with Geneious Prime software version 2020.2.5 (Biomatters Ltd., Auckland, New Zealand) and the consensus sequence between the forward and reverse sequences was extracted for further analyses. Visualization of alignment of the 3 sequencing products to smallRNA-1 was also performed in Geneious Prime software.

**Supplementary Results and Discussion**

A total of 252 tRNAs were predicted in the Nf69 genome. We filtered the results for scores >55 as recommended for functional tRNAs and this left 204 predicted tRNAs. The tRNA types are enumerated in Supplementary Table S8 and the three most prevalent were Met, Lys, and Glu. Upon comparison of percentages of tRNA types to previously predicted tRNA genes in *N. gruberi*[8], we found that for both *Nf* and *N. gruberi*, the three least abundant tRNA genes were Cys, His, and Trp. Accordingly, the two most abundant tRNA genes between the two species were Glu and Lys. Differences between the two species range from a higher prevalence of Ala, Leu, and Ser in *N. gruberi* compared to *Nf*, and a lower prevalence of Met in *N. gruberi* compared to *Nf* (Table S8).

We confirmed the presence of 13 predicted tRNAs in the *Nf* EV sequencing results (Table S1). When comparing the predicted smallRNAs within the EV sequencing results to the tRNA predictions, we noted a phenomenon that we believe is indicative of processing of tRNAs to tRNA fragments (tRFs) as indicated by the overlap of smaller RNAs stacking onto the same scaffold locations that tRNAs were predicted with tRNAscan-SE (Table S1). Interestingly, this is the first report of tRNA editing of any sort in *Nf*. Of the 5 tRFs that we identified, 3 originated from glutamine tRNAs and 2 originated from asparagine tRNAs. For three of the tRFs, tRNA cleavage occurred in the D-loop, and 2 were cleaved in the T-loop. This phenomenon of processing tRNAs into tRNA fragments or halves has been recently reported in another pathogenic amoeba, *Entamoeba histolytica* as a response to stress[9]. Further investigation of this process in *Nf* is warranted to uncover the mechanistic implications behind this processing.
In determining the type of small RNA that the most prevalent smallRNA-1 is, we performed a nucleotide BLAST search and returned hits fell within the 18S ribosomal RNA gene of various isolates of *N. fowleri*. Upon further analysis of the gene region with Integrated Genome Viewer, we identified an ~5.9kb region in Nf69 scaffold 311 of ribosomal RNA (rRNA) with *N. fowleri* EV small RNA reads stacking at various points along the genomic region (Supplemental Figure 8). Additionally, smallRNA-1 and smallRNA-4 were identified and are encoded within this rRNA region—indicating that there is rRNA editing occurring allowing for the packaging and secretion of rRNA fragments within *N. fowleri* EVs. It is possible that the secretion of these small RNAs could allow the amoebae to participate in intracellular communication, signaling, or quorum sensing among amoebae populations and/or other species found within the niches that the amoebae colonize.

### Supplementary Tables

**Table S1.** Identified small RNAs, microRNAs, tRNAs and tRNA fragments.

| Identity | Type | Length (bp) | Sequence | Avg. # of reads in EV seq. | qPCR conf. | Location (scaffold: bp range) | Notes |
|---|---|---|---|---|---|---|---|
| smallRNA-1 | small RNA | 34 | 5'-GGUCCUCGUCUGGCAGUUGCCUCGUAUUCACUCG-3' | 852,500 | Y | 311: 63,160-63,193 | |
| smallRNA-2 | small RNA | 29 | 5'-UUCCAGACUGUCACUCUGGCGACCCGGGU-3' | 253,500 | Y | 69: 24,551-24,579 | |
| smallRNA-3 | small RNA | 42 | 5'-UGGCACCUGCUCACUUAUGCUCCGUUGACGGUCUGCUCCUGG-3' | 667,000 | N | 28: 138,420-138,461 | |
| smallRNA-4 | small RNA | 59 | 5'-UCCAUUAAAGUGCUAAGAGUCUAGUUUCAUAUCGUAGGAGUGGGAUGUUGAGGUUCCAU-3' | 366,500 | N | 311: 61,441-61,499 | |
| smallRNA-5 | tRNA fragment | 53 | 5'-UGGGCCUAGAGUUCCCACUCUCGGAACACAGGAUUGGUAAUCUGAUGUCGCCU-3' | 285,000 | N | 68: 284,768-284,820 | tRNA fragment of s68.tRNA2-GluCTC |
| smallRNA-6 | small RNA | 53 | 5'-AGGCGACAUCAGAUUACCAAUCCUGUGUUCCGAGAGUGGGAACUCUAGGCCCA-3' | 352,000 | N | 49: 18,967-19,019 | |
| smallRNA-7 | tRNA fragment | 52 | 5'-AGCCCACACCAUAUCACCAUCGUAAGGUCUGACAGUGAGACCGCUGGGCCCA-3' | 260,000 | N | 960: 6,983-7,034 | tRNA fragment of s960.tRNA1-AspGTC |
| smallRNA-8 | tRNA fragment | 53 | 5'-UGGGCCUAGAGUUCCCACUCUCGGAACACAGGAUUGGUAAUCUGAUGUCGCCU-3' | 350,000 | N | 68: 283,046-283,098 | tRNA fragment of s68.tRNA3-GluCTC |
| smallRNA-9 | tRNA fragment | 53 | 5'-AGGCGACAUCAGAUUACCAAUCCUGUGUUCCGAGAGUGGGAACUCUAGGCCCA-3' | 355,000 | N | 13: 6,925-6,977 | tRNA fragment of s13.tRNA1-GluCTC |
| smallRNA-10 | tRNA fragment | 52 | 5'-UGGGCCCAGCGGUCUCACUGUCAGACCUUACGAUGGUGAUAUGGUGUGGGCU-3' | 253,000 | N | 64: 93,520-93571 | tRNA fragment of s64.tRNA5-AspGTC |
| nfo-mir-1/5s | microRNA | 23 | 5'-UUCAUUCAACAUUGAUUCAGACU-3' | 1078 | Y | 3 locations on 26 (+): 128,661-128,738; 130,539-130616; 132,029-132106 * | |
| nfo-mir-2 | microRNA | 21 | 5'-UGGUUGUCAUGGUAGAGUGCC-3' | N/A | Y | 135 (-): 50,859-50,935 * | |
| nfo-mir-3 | microRNA | 23 | 5'-UCGAGUAACAUAUGGAAAUGUGU-3' | N/A | Y | 33 (-):180,924-180,999 * | |
| nfo-mir-4/10s | microRNA | 23 | 5'-UUUGUUAAAAUGCGAGACACAGA-3' | 17.5 | N | 110 (+): 281,314-281,401 * | |

| | | | | | | | |
|---|---|---|---|---|---|---|---|
| nfo-mir-5 | microRNA | 23 | 5'-CCUAACCAUUUCUUCUUUCCAGC-3' | N/A | N | 56 (+): 22,981-23,051 * | |
| nfo-mir-6 | microRNA | 22 | 5'-CAUGAACACAUGGACAACUCAU-3' | N/A | N | 378 (+):1,231-1,309 * | |
| nfo-mir-7 | microRNA | 22 | 5'-UUAGCGCGAUGAAAUUUAGGGA-3' | N/A | N | 150 (+):43,158-43,233 * | |
| nfo-mir-8 | microRNA | 21 | 5'-UAUAAAGGCUUCAUUUUCCCA-3' | N/A | N | 250 (+): 17,559-17,630 * | |
| nfo-mir-9 | microRNA | 23 | 5'-UGGUCUUCCUCUUUUUUUCUUGC-3' | N/A | N | 139 (-): 24,452-24,521 * | |
| nfo-mir-10 | microRNA | 24 | 5'-UGCUGCAUGAAAUAAGUAUUCUGU-3' | N/A | N | 2 (+): 203,998-204,057 * | |
| nfo-mir-11 | microRNA | 25 | 5'-UGGGAAAAUGAAGCCUUUAUACGUA-3' | N/A | N | 250 (-): 17,557-17,630 * | |
| nfo-mir-12/9s | microRNA | 23 | 5'-UAGGAGGUUUGUAACAAAUAGAC-3' | 55 | N | 6 (+): 193,964-194,041 * | |
| nfo-mir-1s | microRNA | 24 | 5'-GCUCCCUCCAACCAUCUCCUGAUG-3' | 4041.5 | N | 311 (+): 63,218-63,269 * | |
| nfo-mir-2s | microRNA | 24 | 5'-AAGGUGUGUGGUUAUAAAAUUAUC-3' | 2658 | N | 119 (-/+): 65,151-65,235 * | |
| nfo-mir-3s | microRNA | 23 | 5'-CUAUGCAAGAACAACGAACUCGU-3' | 2161 | N | 2 locations on 69: (+) 61,580-61,646; (-) 63,205-63,271 * | |
| nfo-mir-4s | microRNA | 23 | 5'-ACGAGUUCGUUGUUCUUGCAUAG-3' | 2053.5 | N | 3 locations on 69: (-) 61,580-61,646; (+) 63,205-63,271 * | |
| nfo-mir-6s | microRNA | 23 | 5'-AGUCUGAAUCAAUGUUGAAUGAA-3' | 1012 | N | 3 locations on 26 (-): 128,661-128,735; 130,539-130613; 132,029-132,103 * | |
| nfo-mir-7s | microRNA | 23 | 5'-CACAUUUCCACGUGUUACUCGAC-3' | 573 | N | 33 (-): 180,925-180,999 * | |
| nfo-mir-8s | microRNA | 25 | 5'-UCGGAAAAUGGAGGCCUUUGUACGU-3' | 103.5 | N | 250 (+): 17,559-17,633 * | only in EV S2 |
| s68.tRNA3-GluCTC | tRNA | 74 | 5'-UCGAGGCGACGGCCCUUAGCUUGGGCCUAGAGUUCCCACUCUCGGAACACAGGAUUGGUAAUCUGAUGUCGCCU-3' | 350,000 | Y | 68: 283,027-283,098 | processed into tRNA fragment (smallRNA-8); Score: 67.5 |
| s68.tRNA2-GluCTC | tRNA | 73 | 5'-CGAGGCGACGGCCCUUAGCUUGGGCCUAGAGUUCCCACUCUCGGAACACAGGAUUGGUAAUCUGAUGUCGCCU-3' | 325,000 | N | 68: 284,749-284,820 | processed into tRNA fragment (smallRNA-5); Score: 67.5 |
| s26.tRNA1-AspGTC | tRNA | 70 | 5'-AGCCCACACCAUAUCACCAUCGUAAGGUCUGACAGUGAGACCGCUGGGCCCAAGUUAAGGGCCGUGAGCU-3' | 300,000 | Y | 26: 186,379-186,449 | Score: 58.6 |

| | | | | | | | |
|---|---|---|---|---|---|---|---|
| s162.tRNA1-AspGTC | tRNA | 71 | 5'-GAGCUCACGGCCCUUAACUUGGGCCCAGCGGUCUCACUGUCAGACCUUACGAUGGUGAUAUGGUGUGGGCU-3' | 300,000 | N | 162: 54,358-54,429 | Score: 58.8 |
| s165.tRNA2-AspGTC | tRNA | 71 | 5'-GAGCUCACGGCCCUUAACUUGGGCCCAGCGGUCUCACUGUCAGACCUUACGAUGGUGAUAUGGUGUGGGCU-3' | 300,000 | N | 165: 46,530-46,600 | Score: 58.6 |
| s960.tRNA1-AspGTC | tRNA | 71 | 5'-AGCCCACACCAUAUCACCAUCGUAAGGUCUGACAGUGAGACCGCUGGGCCCAAGUUAAGGGCCGUGAGCUC-3' | 300,000 | N | 960: 6,983-7,053 | processed into tRNA fragment (smallRNA-7); Score: 58.6 |
| s65.tRNA2-AspGTC | tRNA | 71 | 5'-GAGCUCACGGCCCUUAACUUGGGCCCAGCGGUCUCACUGUCAGACCUUACGAUGGUGAUAUGGUGUGGGCU-3' | 275,000 | N | 65: 27,377-27,447 | Score: 58.6 |
| s15.tRNA1-AspGTC | tRNA | 72 | 5'-UUAGCCCACACCAUAUCACCAUCGUAAGGUCUGACAGUGAGACCGCUGGGCCCAAGUUAAGGGCCGUGAGCU-3' | 200,000 | N | 15: 65,817-65,887 | Score: 58.6 |
| s64.tRNA5-AspGTC | tRNA | 71 | 5'-GAGCUCACGGCCCUUAACUUGGGCCCAGCGGUCUCACUGUCAGACCUUACGAUGGUGAUAUGGUGUGGGCU-3' | 100,000 | N | 64: 93,500-93,571 | processed into tRNA fragment (smallRNA-10); Score: 58.8 |
| s13.tRNA1-GluCTC | tRNA | 72 | 5'-AGGCGACAUCAGAUUACCAAUCCUGUGUUCCGAGAGUGGGAACUCUAGGCCCAAGCUAAGGGCCGUCGCCUC-3' | 30,000 | N | 13: 6,925-6,996 | processed into tRNA fragment (smallRNA-9); Score: 67.5 |
| s196.tRNA1-HisGTG | tRNA | 71 | 5'-AUCCUCGUCUCCCUUAGCUUGGGAGCAGAUUCCUUGGUGUUAGGAAGUAAGAUGGUAACUUGAUACGGGGA-3' | 10,000 | N | 196: 13,761-13,831 | Score: 65.5 |
| s14.tRNA1-HisGTG | tRNA | 71 | 5'-AUCCUCGUCUCCCUUAGCUUGGGAGCAGAUUCCUUGGUGUUAGGAAGUAAGAUGGUAACUUGAUACGGGGA-3' | 7,000 | N | 14: 7,407-7,477 | Score: 65.5 |
| s41.tRNA1-HisGTG | tRNA | 72 | 5'-AAUCCUCGUCUCCCUUAGCUUGGGAGCAGAUUCCUUGGUGUUAGGAAGUAAGAUGGUAACUUGAUACGGGGA-3' | 5,000 | N | 41: 31,761-31,831 | Score: 65.5 |

* = location of microRNA precursor coordinates; (+/-) = + or - strand of genome.

We utilized a tRNAscan-SE Score cut off >55.

Like sequences among predictions are highlighted in coordinating colors with mismatching basepairs colored red.

**Table S2.** Taqman assay information.

| target | assay ID | cat# |
|---|---|---|
| cel-mir-39 | 000200 | 4440887 |
| smallRNA-1 | CTPRKJ6 | 4398989 |
| smallRNA-2 | CTNKR42 | 4398987 |
| s68.tRNA3-GluCTC | CTH6AAG | 4440418 |
| s26.tRNA1-AspGTC | CTKA3VE | 4440418 |

**Table S3.** Taqman RT-qPCR assay thermal cycling parameters.

| Step | Temp. | Time | # of cycles |
|---|---|---|---|
| Pre-processing | 85°C | 5 min | 1 |
| | 60°C | 5 min | 1 |
| | Place on ice | | |
| Reverse Transcription | 16°C | 30 min | 1 |
| | 42°C | 30 min | 1 |
| | 85°C | 5 min | 1 |
| | 4°C | Hold | |
| qPCR | 50°C | 2 min | 1 |
| | 95°C | 2 min | 1 |
| | 95°C | 1 sec | 45 |
| | 60°C | 20 sec | |

**Table S4.** Conditions for extraction of EVs from amoeba-conditioned media or FBS with isolation reagent.

| Contents | Media | EV-dep. FBS? | # of cells seeded | Final cell conc. | Culture vol. (mL) | Culture time (h) | Flask Size ($cm^2$) | Purpose |
|---|---|---|---|---|---|---|---|---|
| BP-Nf69 Pass 0 rep. 1 | NCM | Y | from brain | 33,625,000 | 100 | 216 | 175 | small RNA sequencing |
| BP-Nf69 Pass 1 rep. 2 | NCM | Y | 6,000,000 | 32,625,000 | 100 | 77 | 175 | small RNA sequencing |
| Nf69 | NCM | N | 750,000 | 14,500,000 | 25 | 90.5 | 75 | smallRNA-1 detection |
| *N. lovaniensis* | NCM | N | 750,000 | 7,680,000 | 25 | 114.5 | 75 | smallRNA-1 detection |
| *N. gruberi* | M7 | N | 750,000 | 8,625,000 | 25 | 68.5 | 75 | smallRNA-1 detection |
| *A. castellanii* | PG | n/a | 500,000 | 17,380,000 | 25 | 90.5 | 75 | smallRNA-1 detection |
| *B. mandrillaris* | BMI | N | 750,000 * | 8,750,000 | 3x 8.33 | 90.5 | 3x 75 | smallRNA-1 detection |
| FBS (4 replicates) | FBS | N | n/a | n/a | 2.5 | n/a | n/a | control |
| * = seeded into 3 flasks and combined after incubation due to sensitivity to high confluency in culture. | | | | | | | | |

**Table S5.** Details and assay results from all control/uninfected human biofluids tested.

| Name | Fluid | Volume tested (μL) | Assay | Pooled? | Source | mean smallRNA-1 Cq | mean smallRNA-2 Cq |
|---|---|---|---|---|---|---|---|
| CSF-1 | CSF | 500 | smallRNA-1 | Y | AdventHealth | no signal | n/a |
| CSF-2 | CSF | 500 | smallRNA-1 | Y | AdventHealth | 36.4 | n/a |
| CSF-3 | CSF | 500 | smallRNA-1 | Y | AdventHealth | 35.8 | n/a |
| CDC CSF-11 | CSF | 15 | smallRNA-1 | N | CDC | no signal | n/a |
| CDC CSF-12 | CSF | 15 | smallRNA-1 | N | CDC | no signal | n/a |
| 237/F/Black/54yo | plasma | 250 | smallRNA-1 and -2 | N | Grifols | 40.2 | 39.2 |
| 177/F/White/23yo | plasma | 100 | smallRNA-1 and -2 | N | Grifols | 39.2 | no signal |
| 238/M/Black/29yo | plasma | 100 | smallRNA-1 | N | Grifols | 41.8 | n/a |
| 207/M/Black/32yo | plasma | 250 | smallRNA-1 | N | Grifols | no signal | n/a |
| 229/M/Latino/51yo | plasma | 225 | smallRNA-1 | N | Grifols | 39.8 | n/a |
| 234/F/Black/53yo | plasma | 200 | smallRNA-1 | N | Grifols | 42.6 | n/a |
| 388/M/Black/51yo | plasma | 200 | smallRNA-1 | N | Grifols | 37.1 | n/a |
| Plasma-1 | plasma | 500 | smallRNA-1 | Y | AdventHealth | 41.8 | n/a |
| Plasma-2 | plasma | 500 | smallRNA-1 | Y | AdventHealth | no signal | n/a |
| Plasma-3 | plasma | 500 | smallRNA-1 | Y | AdventHealth | no signal | n/a |
| Serum-1 | serum | 500 | smallRNA-1 | Y | AdventHealth | no signal | n/a |
| Serum-2 | serum | 500 | smallRNA-1 | Y | AdventHealth | 35.7 | n/a |
| Serum-3 | serum | 500 | smallRNA-1 | Y | AdventHealth | no signal | n/a |
| CDC Serum S2 | serum | 20 | smallRNA-1 | N | CDC | no signal | n/a |
| CDC Serum S3 | serum | 20 | smallRNA-1 | N | CDC | 39.8 | n/a |
| CDC Serum S4 | serum | 20 | smallRNA-1 | N | CDC | 38.7 | n/a |
| Urine-1 | urine | 500 | smallRNA-1 | N | AdventHealth | 42.5 | n/a |
| Urine-2 | urine | 500 | smallRNA-1 | N | AdventHealth | 33.4 | n/a |
| Urine-3 | urine | 500 | smallRNA-1 | N | AdventHealth | no signal | n/a |
| Urine-4 | urine | 500 | smallRNA-1 | N | AdventHealth | 35 | n/a |
| Urine-5 | urine | 500 | smallRNA-1 | N | AdventHealth | no signal | n/a |
| Urine-6 | urine | 500 | smallRNA-1 | N | AdventHealth | 34.8 | n/a |
| Urine-7 | urine | 500 | smallRNA-1 | N | AdventHealth | 33.7 | n/a |
| Urine-8 | urine | 500 | smallRNA-1 | N | AdventHealth | 38.5 | n/a |

**Table S6.** microRNA validation primer sequences and annealing temperatures.

| Assay | Target | Primer | Sequence | Conc. Used | Anneal temp. |
|---|---|---|---|---|---|
| Reverse Transcription | nfo-mir-1 | stem-loop | 5'-GTCGTATCCAGTGCAGGGTCCGAGGTATTCGCACTGGATACGACCATCAG-3' | 375 pM | - |
| | nfo-mir-2 | stem-loop | 5'-GTCGTATCCAGTGCAGGGTCCGAGGTATTCGCACTGGATACGACGATAAT-3' | 375 pM | - |
| | nfo-mir-3 | stem-loop | 5'-GTCGTATCCAGTGCAGGGTCCGAGGTATTCGCACTGGATACGACACACAT-3' | 375 pM | - |
| SYBR green qPCR | universal | Reverse | 5'-CCAGTGCAGGGTCCGAGGTA-3' | 500 nM | - |
| | nfo-mir-1 | Forward | 5'-TACTCAGCTCCCTCCAACCATCTC-3' | 500 nM | 54°C |
| | nfo-mir-2 | Forward | 5'-GCAGGCTTGGTTGTCATGGTAGAG-3' | 500 nM | 60°C |
| | nfo-mir-3 | Forward | 5'-GTGGCAGGATCGAGTAACATATGGAAAT-3' | 500 nM | 56°C |

**Table S7.** SYBR green RT-qPCR assay thermal cycling parameters.

| Step | Temp. | Time | # of cycles |
|---|---|---|---|
| Reverse Transcription | 16°C | 30 min | 1 |
| | 42°C | 30 min | 1 |
| | 85°C | 5 min | 1 |
| | 4°C | Hold | |
| qPCR | 95°C | 2 min | 1 |
| | 95°C | 15 sec | 40 |
| | Variable | 1 min | |

**Table S8.** Enumeration of tRNA types reported in the literature for *N. gruberi* (872 total tRNA genes) compared to the determinations made for *Nf* in this study using tRNAscan-SE (204 total tRNA genes). Cells with percentages of total are color-coded according to tRNA gene abundance.

| tRNA type | *N. gruberi* count | *N. fowleri* count | *N. gruberi* % | *N. fowleri* % |
|---|---|---|---|---|
| Ala | 49 | 5 | 5.62 | 2.45 |
| Arg | 48 | 12 | 5.50 | 5.88 |
| Asn | 34 | 9 | 3.90 | 4.41 |
| Asp | 52 | 10 | 5.96 | 4.90 |
| Cys | 18 | 5 | 2.06 | 2.45 |
| Gln | 38 | 11 | 4.36 | 5.39 |
| Glu | 69 | 14 | 7.91 | 6.86 |
| Gly | 56 | 15 | 6.42 | 7.35 |
| His | 21 | 5 | 2.41 | 2.45 |
| Ile | 45 | 12 | 5.16 | 5.88 |
| Leu | 75 | 10 | 8.60 | 4.90 |
| Lys | 74 | 16 | 8.49 | 7.84 |
| Met | 39 | 17 | 4.47 | 8.33 |
| Phe | 28 | 6 | 3.21 | 2.94 |
| Pro | 31 | 10 | 3.56 | 4.90 |
| Ser | 61 | 11 | 7.00 | 5.39 |
| Thr | 47 | 12 | 5.39 | 5.88 |
| Trp | 16 | 5 | 1.83 | 2.45 |
| Tyr | 28 | 8 | 3.21 | 3.92 |
| Val | 43 | 10 | 4.93 | 4.90 |

gene abundance:

- higher
- lower

**Table S9:** Description of clinical samples from CDC and previous testing results.

| Information Provided by Centers for Disease Control and Prevention | | | | | | Current Study |
|---|---|---|---|---|---|---|
| Specimen type | Date of collection | Results | Assay performed | Comments | FLA real-time PCR CT values † | smallRNA-1 RT-qPCR mean Cq values * |
| CSF | 8/2022 | Positive for *Nf* | FLA real-time PCR | -- | *Nf*: 29·05 | 26 |
| CSF | 7/2022 | Positive for *Nf* | FLA real-time PCR | -- | *Nf*: 26·10 | 26..5 |
| CSF | 9/2021 | Positive for *Nf* | FLA real-time PCR | -- | *Nf*: 28·06 | 24.3 |
| CSF | 8/2021 | Positive for *Nf* | FLA real-time PCR | -- | *Nf*: 26·40 | 30.5 |
| CSF | 8/2021 | Positive for *Nf* | FLA real-time PCR | -- | *Nf*: 27·20 | 28 |
| CSF | 9/2020 | Positive for *Nf* | FLA real-time PCR | -- | *Nf*: 22·10 | 24 |
| CSF | 2/2023 | Positive for *Ac* | FLA real-time PCR | -- | *Ac* : 29·10 | no signal |
| CSF | 12/2022 | Positive for *Ac* | FLA real-time PCR | -- | *Ac* : 35·80 | no signal |
| CSF | 8/2021 | Positive for *Bm* | FLA real-time PCR | -- | *Bm* : 36·90 | no signal |
| CSF | 2/2021 | Positive for *Bm* | FLA real-time PCR | -- | *Bm* : 32·80 | no signal |
| CSF | 11/2022 | Negative | FLA real-time PCR | -- | no signal | no signal |
| CSF | 8/2022 | Negative | FLA real-time PCR | -- | no signal | no signal |
| Whole Blood | 9/7/16 | Positive for *Nf* (weak) | FLA real-time PCR | These samples were from the same PAM patient and may have been exposed to room temp. for days/weeks before moved to -20ºC or lower. | *Nf*: 37.4 | 32.4 |
| Plasma | 9/7/16 | Positive for *Nf* (very weak) | FLA real-time PCR | | *Nf*: 38.2 | no signal |
| Whole Blood | 2/18/20 | Positive for *Nf* | FLA real-time PCR | Another PAM patient. | *Nf*: 36.4 | 32.6 |
| Serum | 2/16/22 | Negative | IFA serology assay | Negative control samples. Only serological test, IFA was performed on these samples. FLA real-time PCR was not performed. | N/A | 41.8 |
| Serum | 11/16/21 | Negative | IFA serology assay | | N/A | no signal |
| Serum | 8/26/21 | Negative | IFA serology assay | | N/A | 39.8 |
| Serum | 8/19/21 | Negative | IFA serology assay | | N/A | 38.7 |
| Serum | 1/14/21 | Positive for anti-*Bm* Ab | IFA @1:128 titer | | N/A | no signal |
| Serum | 2/8/18 | Positive for anti-*Bm* Ab | IFA @1:128 titer | | N/A | no signal |
| Serum | 9/14/16 | Positive for anti-*Bm* Ab | IFA @1:64 titer | | N/A | no signal |
| Serum | 11/14/16 | Positive for anti-*Ac* Ab | IFA @ 1:64 titer | | N/A | no signal |
| Serum | 8/11/16 | Negative | IFA serology assay | Serum samples from the same recovered PAM patient, but only the serum #11 was seropositive for *Nf*. These were not tested for *Nf* DNA by FLA real-time PCR. | N/A | 39.1 |
| Serum | 6/16/16 | Negative | IFA serology assay | | N/A | no signal |
| Serum | 8/23/16 | Positive for anti-*Nf* Ab | IFA @1:128 | | N/A | 41.8 |

N/A = Not applicable; Ab = Antibody; *Nf* = *Naegleria fowleri* ; *Bm* = *Balamuthia mandrillaris* ; *Ac* = *Acanthamoeba* spp.

† = For the FLA real-time PCR, an input volume of 200µL is used; a CT value of >37 (but <40) is considered weak positive.

* = For the smallRNA-1 assay, qPCR is run for 45 cycles; the positivity cut-off for human biofluids is a mean Cq of <33·4 among the 3 tech. replicates.

**A**

| Whole Cell Extract Predictions | | | EV-seq |
|---|---|---|---|
| **Putative Name** | **Mature miRNA sequence** | **Predicted Stem-loop Structure** | **Confirmation** |
| **nfo-mir-1** | 5'-UUCAUUCAACAUUGAUUCAGACU-3' | 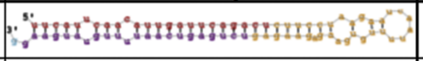 | **Yes** |
| **nfo-mir-2** | 5'-UGGUUGUCAUGGUAGAGUGCC-3' | 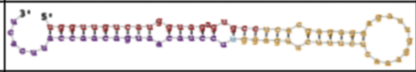 | **Maybe** -1 of 2 samples |
| **nfo-mir-3** | 5'-UCGAGUAACAUAUGGAAAUGUGU-3' | 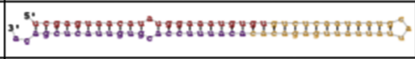 | **Yes** -Star Sequence |
| **nfo-mir-4** | 5'-UUUGUUAAAAUGCGAGACACAGA-3' | 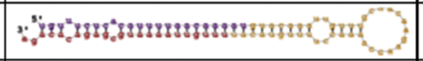 | **Yes** -Lower Prevalence |
| **nfo-mir-5** | 5'-CCUAACCAUUUCUUCUUUCCAGC-3' | 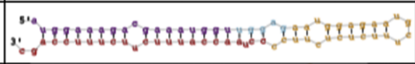 | **Maybe** -1 of 2 samples |
| **nfo-mir-6** | 5'-CAUGAACACAUGGACAACUCAU-3' | 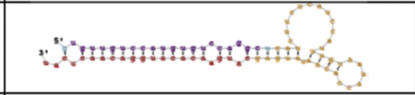 | **No** |
| **nfo-mir-7** | 5'-UUAGCGCGAUGAAAUUUAGGGA-3' | 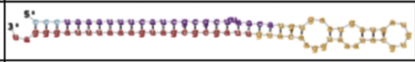 | **Maybe** -1 of 2 samples -Star Sequence |
| **nfo-mir-8** | 5'-UAUAAAGGCUUCAUUUUCCCA-3' | 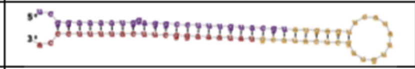 | **Yes** -Star Sequence |
| **nfo-mir-9** | 5'-UGGUCUUCCUCUUUUUUUCUUGC-3' | 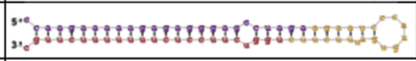 | **Maybe** -1 of 2 samples |
| **nfo-mir-10** | 5'-UGCUGCAUGAAAUAAGUAUUCUGU-3' | 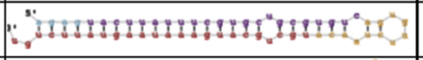 | **No** |
| **nfo-mir-11** | 5'-UGGGAAAAUGAAGCCUUUAUACGUA-3' | 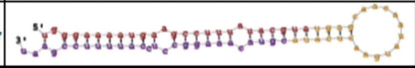 | **Maybe** -1 of 2 samples -Star Sequence |
| **nfo-mir-12** | 5'-UAGGAGGUUUGUAACAAAUAGAC-3' | 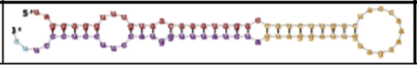 | **Yes** |

**B**

| EV Extract Predictions | | | |
|---|---|---|---|
| **Putative Name** | **Mature miRNA sequence** | **Predicted Stem-loop Structure** | **Avg. # of Reads** |
| **nfo-mir-1s** | 5'-GCUCCCUCCAACCAUCUCCUGAUG-3' | 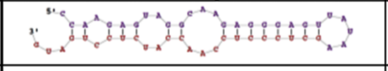 | 4041.5 |
| **nfo-mir-2s** | 5'-AAGGUGUGUGGUUAUAAAAUUAUC-3' | 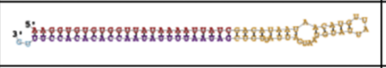 | 2658 |
| **nfo-mir-3s** | 5'-CUAUGCAAGAACAACGAACUCGU-3' | 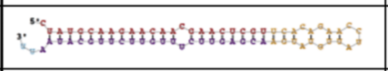 | 2161 |
| **nfo-mir-4s** | 5'-ACGAGUUCGUUGUUCUUGCAUAG-3' | 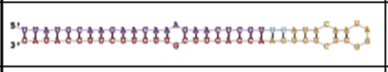 | 2053.5 |
| **nfo-mir-5s** (Previously nfo-mir-1) | 5'-UUCAUUCAACAUUGAUUCAGACU-3' | 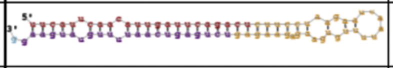 | 1078 |
| **nfo-mir-6s** | 5'-AGUCUGAAUCAAUGUUGAAUGAA-3' | 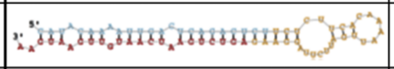 | 1012 |
| **nfo-mir-7s** (Previously nfo-mir-3*) | 5'-CACAUUUCCACGUGUUACUCGAC-3' | 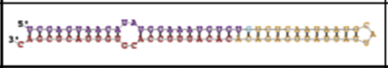 | 573 |
| **nfo-mir-8s** (Previously nfo-mir-8*) | 5'-UCGGAAAAUGGAGGCCUUUGUACGU-3' | 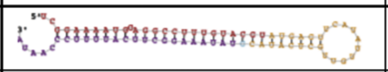 | 103.5 |
| **nfo-mir-9s** (Previously nfo-mir-12) | 5'-UAGGAGGUUUGUAACAAAUAGAC-3' | 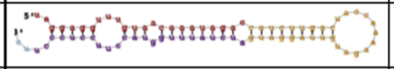 | 55 |
| **nfo-mir-10s** (Previously nfo-mir-4) | 5'-UUUGUUAAAAUGCGAGACACAGA-3' | 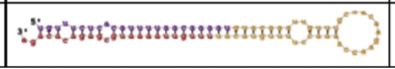 | 17.5 |

**Supplemental Figure 1: miRDeep2 microRNAs predicted in whole cell small RNA sequencing versus EV small RNA sequencing.** (A) The mature microRNA sequence and the predicted stem-loop structure for top 12 most prevalent microRNAs predicted in both replicates of the whole cell small RNA sequencing. These predictions were cross-referenced to results obtained for EV sequencing microRNA prediction. (B) Predicted mature microRNAs and stem-loop structures present in both small RNA *Nf*-EV sequencing samples.

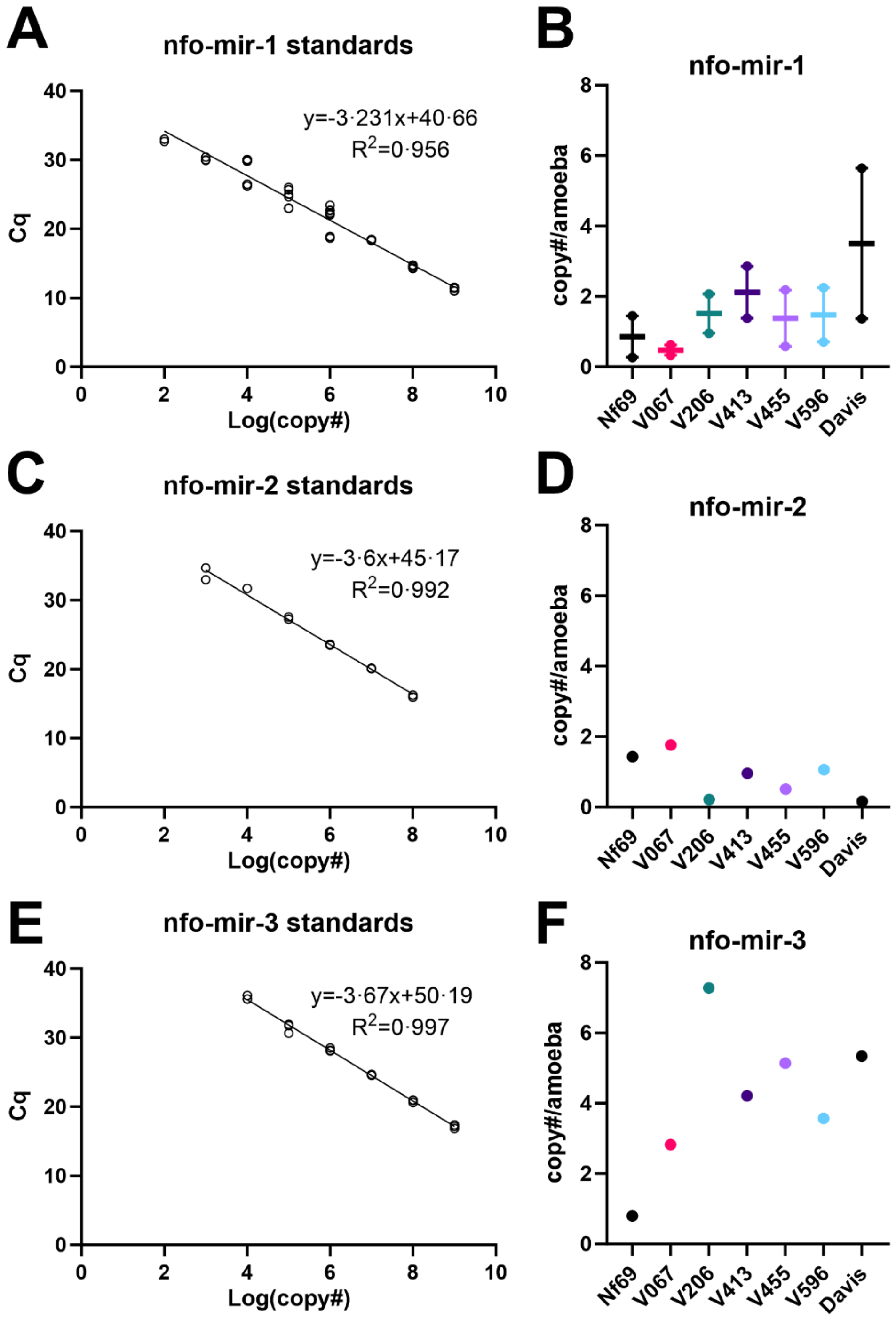

**Supplemental Figure 2: RT-qPCR confirmation of presence of top 3 most prevalent microRNAs in whole amoebae.** (A-B) nfo-mir-1 standard curve and presence in 7 clinical isolates with Cq values ranging from 21.4 to 25.8, (C-D) nfo-mir-2 with Cq values ranging from 25.1 to 29.7, and (E-F) nfo-mir-3 with Cq values ranging from 28 to 30.8. Each data point in panels B, D, and F represents 3 technical replicates in RT-qPCR assay.

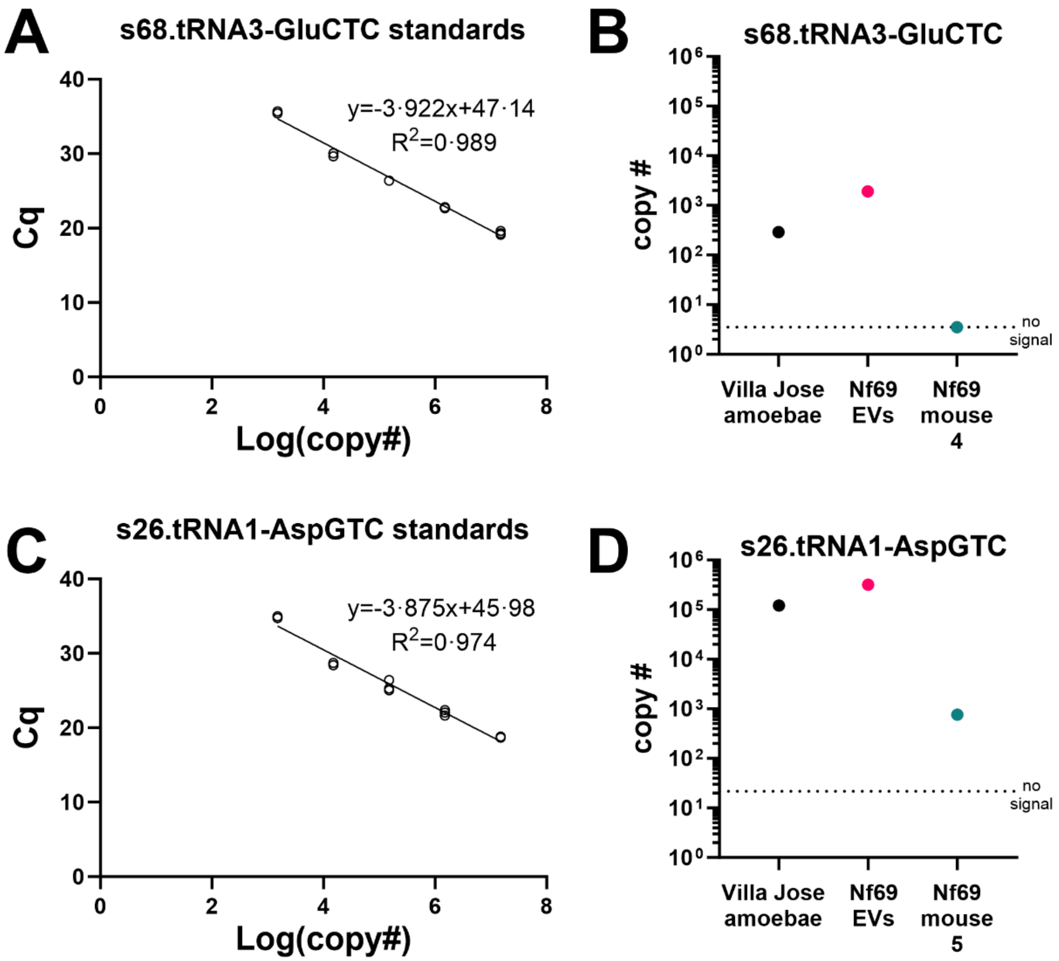

**Supplemental Figure 3: RT-qPCR confirmation of presence of 2 most prevalent tRNAs in *Nf-EVs* and amoebae pellets.** (A) Standard curve of most prevalent tRNA, s68.tRNA3-GluCTC. (B) Detection of s68.tRNA3-GluCTC in Villa Jose amoebae pellet and Nf69 EVs (Cq values from 33.8-37.8), with no signal obtained in the plasma of an end-stage Nf69-infected mouse. (C) Standard curve of third most prevalent tRNA, s26.tRNA1-AspGTC. (D) Detection of s26.tRNA1-AspGTC in Villa Jose amoebae pellet, Nf69 EVs, and the plasma of an end-stage Nf69 infected mouse (Cq values from 24.6-34.9). Each data point in panels B and D represents 3 technical replicates in RT-qPCR assay.

A

B

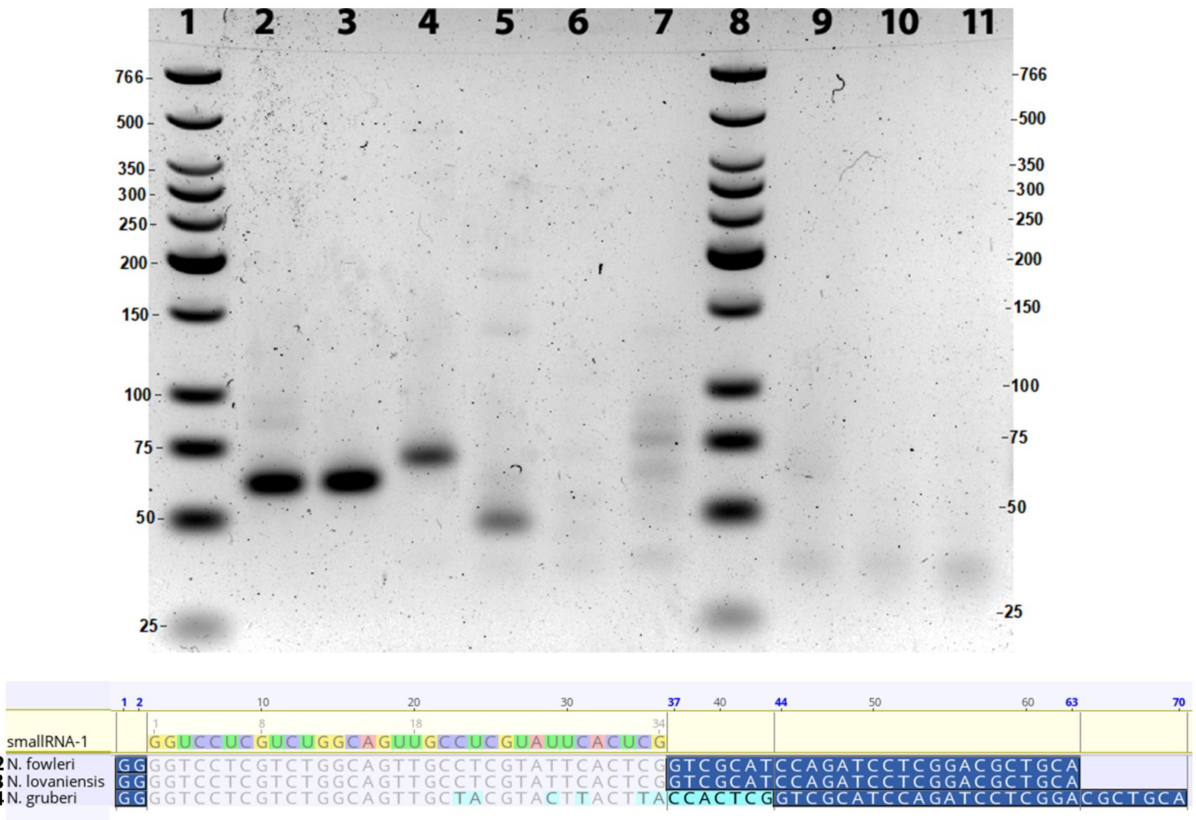

**Supplemental Figure 4: Agarose gel confirmation of qPCR products reported in Figure 2E.** (A) 1-Ladder, 2-*N. fowleri* Nf69 EVs, 3-*N. lovaniensis* EVs, 4-*N. gruberi* EVs, 5-*A. castellanii* EVs, 6-*B. mandrillaris* EVs, 7-fetal bovine serum EVs, 8-Ladder, 9-no template control for reverse transcription, 10-no reverse transcriptase control, 11-no template control for qPCR. (B) Sequence alignment of qPCR products 2, 3 and 4 to smallRNA-1. Dark blue highlighted portions represent primer additions from Custom Taqman assay, light blue highlighted portions in sequence 4 represent mismatches compared to smallRNA-1.

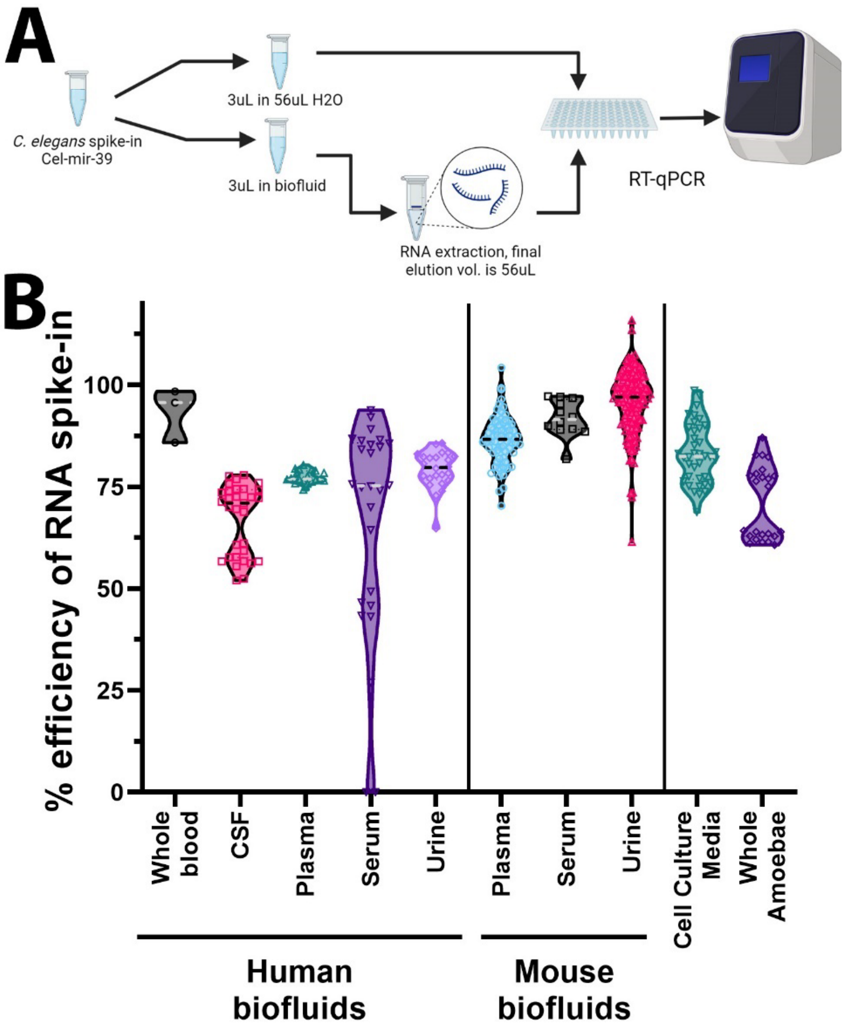

**Supplemental Figure 5: RNA extraction efficiency across sample types tested with RT-qPCR.** (A) Schematic showing process of spike-in of *Caenorhabditis elegans* microRNA cel-mir-39 into biofluids after lysis buffer was added and assayed in parallel to cel-mir-39 diluted in same volume as elution of RNA followed by RT-qPCR to test for RNA extraction efficiency. (B) Percent efficiency of RNA extraction calculated as follows: ((mean control cel-mir-39 in H2O / mean spike-in cel-mir-39) *100), ranged from 0 to >100% as determined by spike-in across human and mouse biofluids as well as cell culture media. Data indicates that human serum inhibits RT-qPCR assay compared to other biofluids. Each data point represents 2 technical replicates in RT-qPCR assay.

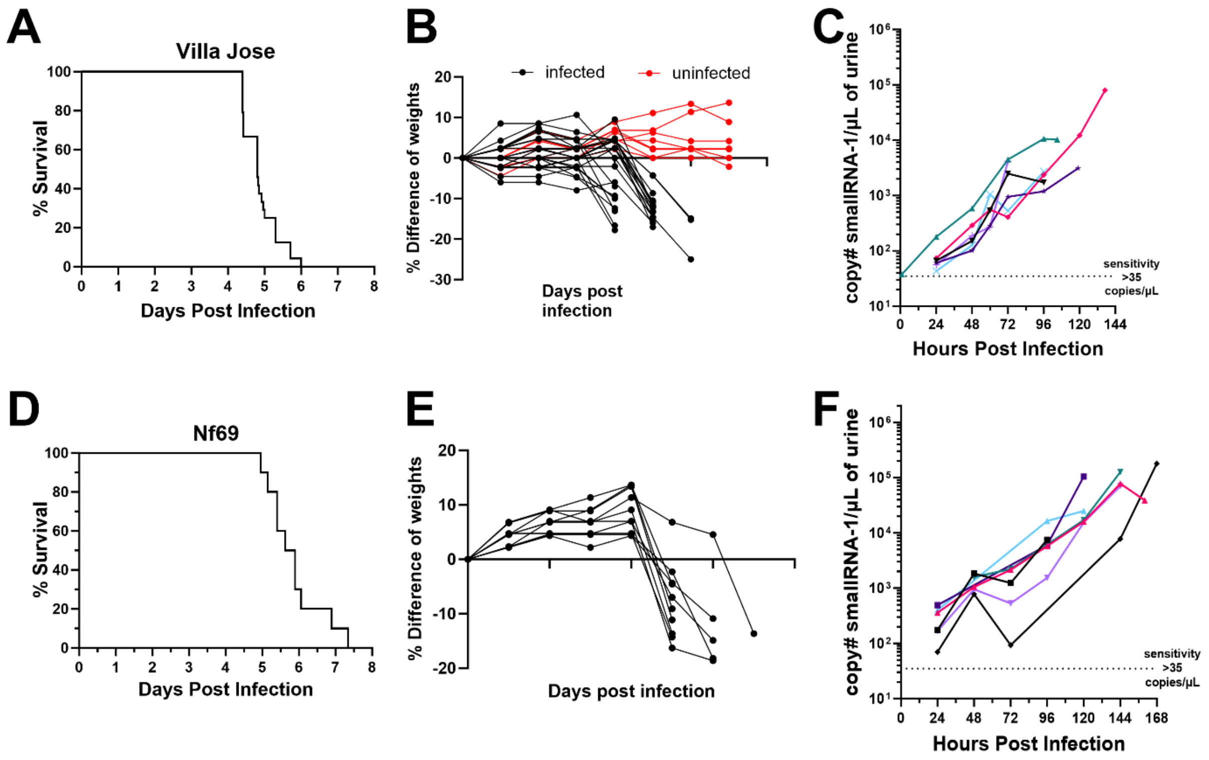

**Supplemental Figure 6: Survival curves, weight tracking and smallRNA-1 tracking in urine of individual mice.** (A) Survival curve of 24 mice infected with 1,000 VJ amoebae from Figure 5B allowed to progress to end-stage of infection. (B) Weight tracking of VJ-infected or uninfected mice shows the rapid decrease in weight associated with PAM infection. (C) SmallRNA-1 tracking over time in the urine of 6 individual mice infected by 1,000 VJ amoebae. (D) Survival curve of 10 mice infected with 5,000 Nf69 amoebae. (E) Weight tracking of Nf69-infected mice. (F) SmallRNA-1 tracking in urine of 6 individual Nf69-infected mice over time.

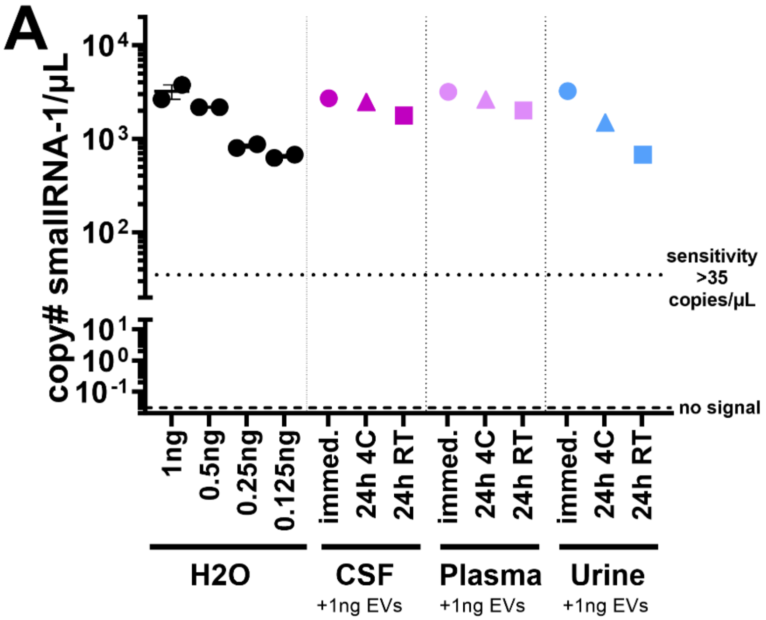

**Supplemental Figure 7: Determination of *Nf*-EV stability and detection of smallRNA-1 in human blood and serum.** (A) Detection of various concentrations of Nf69 EVs spiked into 200µL $H_2O$, and determination of stability of 1ng of EVs spiked into 200µL human CSF, plasma and urine with RNA extracted immediately, after 24h stored at 4°C or after 24h stored at RT. Each data point is representative of 3 technical replicates in RT-qPCR assay.

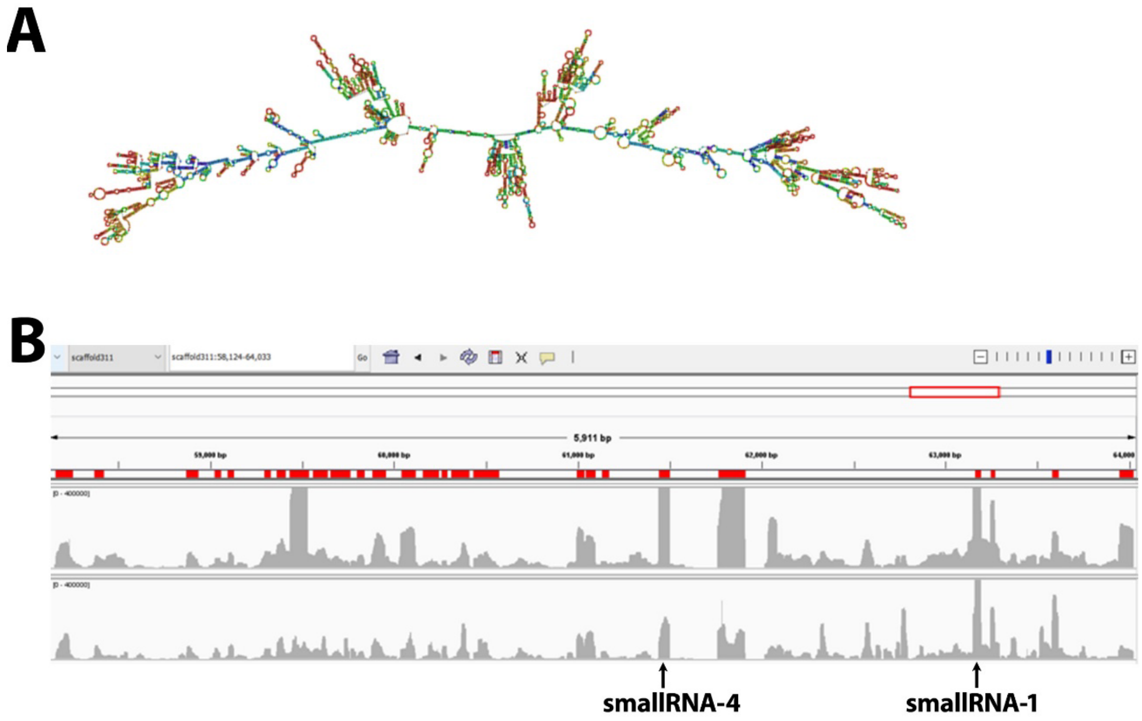

**Supplemental Figure 8: Ribosomal RNA region within scaffold 311 of Nf69 genome.** (A) RNAfold of ~5.9kb region of rRNA. (B) Integrated Genome Viewer view two *Nf*-EV smallRNA sequencing samples aligned to Nf69 genome depicting region of rRNA with annotations for smallRNA-1 and smallRNA-4.
